# High neuron-microglia interaction at the node of Ranvier predicts recovery in an inflammatory model of Multiple Sclerosis

**DOI:** 10.64898/2026.04.14.718431

**Authors:** V. Pantazou, G. Dorcet, T. Roux, M.S Aigrot, C. Perrot, A. Mathias, M. Canales, F.X. Lejeune, B. Stankoff, R. Du Pasquier, C. Lubetzki, A. Desmazières

**Affiliations:** Paris Brain Institute, Sorbonne Université, CNRS, Inserm, GH Pitié-Salpêtrière, Paris, France; Service of neurology, Department of clinical neurosciences, Lausanne University Hospital and University of Lausanne, Lausanne, Switzerland; Neurology Department, Assistance Publique-Hôpitaux de Paris, Pitié Salpêtrière Hospital Paris, France; Laboratory of neuroimmunology, Center of research in neurosciences, Department of clinical neurosciences, Lausanne University Hospital and University of Lausanne, Switzerland; Data Analysis Core, Paris Brain Institute, Paris, France

## Abstract

Microglia, the resident immune cells of the central nervous system, dynamically respond to their environment in health, injury and disease. They contact axons at the nodes of Ranvier in an activity-dependent manner, a process which contributes to repair, but how adaptive immunity in Multiple Sclerosis (MS) impacts this neuron-microglia crosstalk remains unknown. Using an inflammatory MS model, we identify strengthened microglia-node interactions at remission onset, with marked interindividual variability. Increased engagement correlates with a Th2-related cytokine signature, and IL13/IL4 are sufficient to enhance microglia-node contacts. High interaction levels associate with more pro-regenerative microglia, improved tissue repair and better functional recovery. Low-intensity physical exercise at remission onset further promotes microglia-node interaction, pro-regenerative microglia and improves recovery. Consistently, high microglia-node contact in MS tissue is associated with more extensive remyelination, underscoring this interaction as a key process in repair.

## Introduction

Multiple sclerosis (MS) is an inflammatory, demyelinating and neurodegenerative disease of the central nervous system (CNS), affecting over two million people worldwide (1, 2). While, for some patients, endogenous remyelination occurs after demyelination (3), the extent and effectiveness of this process present a high interindividual variability (4, 5). Interestingly, when efficient, remyelination may protect from axonal injury (6–8) and inversely correlates with clinical disability in the first years of the disease (5, 9). It is therefore crucial to decipher the key processes leading to efficient repair in order to limit disability progression in MS.

Immune cells - both adaptive and innate - are major actors of this regenerative process (10–18). As the brain main resident immune cells, microglia can adopt various dynamic states in response to the disruption of brain homeostasis (19–24). Microglial activation is necessary for efficient remyelination, as pro-inflammatory microglia participate in myelin debris phagocytosis as well as recruitment and proliferation of the oligodendrocyte precursor cells (OPCs), while pro-regenerative microglia stimulate oligodendrocyte differentiation and remyelination (11, 18, 25–27). Pro-regenerative microglial cells are in addition involved in the resolution of inflammation and in neuroprotection through secretion of pro-regenerative cytokines and neurotrophic factors (28–30). The timely switch from pro-inflammatory to pro-regenerative microglia appears to be a key factor for efficient repair, and microglia blocked in deleterious pro-inflammatory states are associated with the progression of the disease (31, 32). A deeper understanding of the mechanisms that drive microglia towards pro-regenerative states is therefore essential for promoting repair in MS.

Microglia-neuron interactions occur on various neuronal sub-compartments, including the soma and dendrites, as well as at the axon initial segment and the synapses (33–36). We recently described that a neuron-microglia crosstalk occurs at the nodes of Ranvier in an activity-dependent manner (37, 38). Disrupting this interaction at the onset of repair in a toxic demyelinating model led to a decrease of pro-regenerative microglia and impaired remyelination (37), while a physiological increase of neuronal activity reinforced microglia-node interaction and promoted the switch towards pro-regenerative microglia (38).

Here, we asked in an inflammatory MS model (MOG-EAE) whether adaptive immunity shapes microglia-node interaction and microglial response in repair. At remission onset, microglia-neuron contacts at nodes of Ranvier were markedly increased but showed strong inter-individual variability. High microglia-node interaction aligned with Th2-associated CNS immune signature and associated with more pro-regenerative microglia, enhanced tissue repair, and improved clinical recovery. Low intensity physical exercise at remission onset further amplified these interactions, expanded the pro-regenerative microglial pool and promoted recovery. Consistently, high microglia-node contacts correlated with greater remyelination in MS lesions, strengthening their relevance as a key pro-repair mechanism.

## Results

### Infiltrating macrophages contribute to nodal contacts in the perilesional tissue at disease peak, while microglia-node contacts dominate at remission onset

To study the impact of recruited immunity on microglia-node interaction, we developed a MOG-induced EAE mouse model, a well-characterized inflammatory model of MS with the onset of symptoms occurring in the second week following induction, before reaching a peak followed by a partial remission (Figure 1a-c). As recruited macrophages can contact nodes and initiate perinodal myelin phagocytosis at EAE onset (39), we first asked which innate immune populations contact nodes over the disease course.

**Figure 1.**
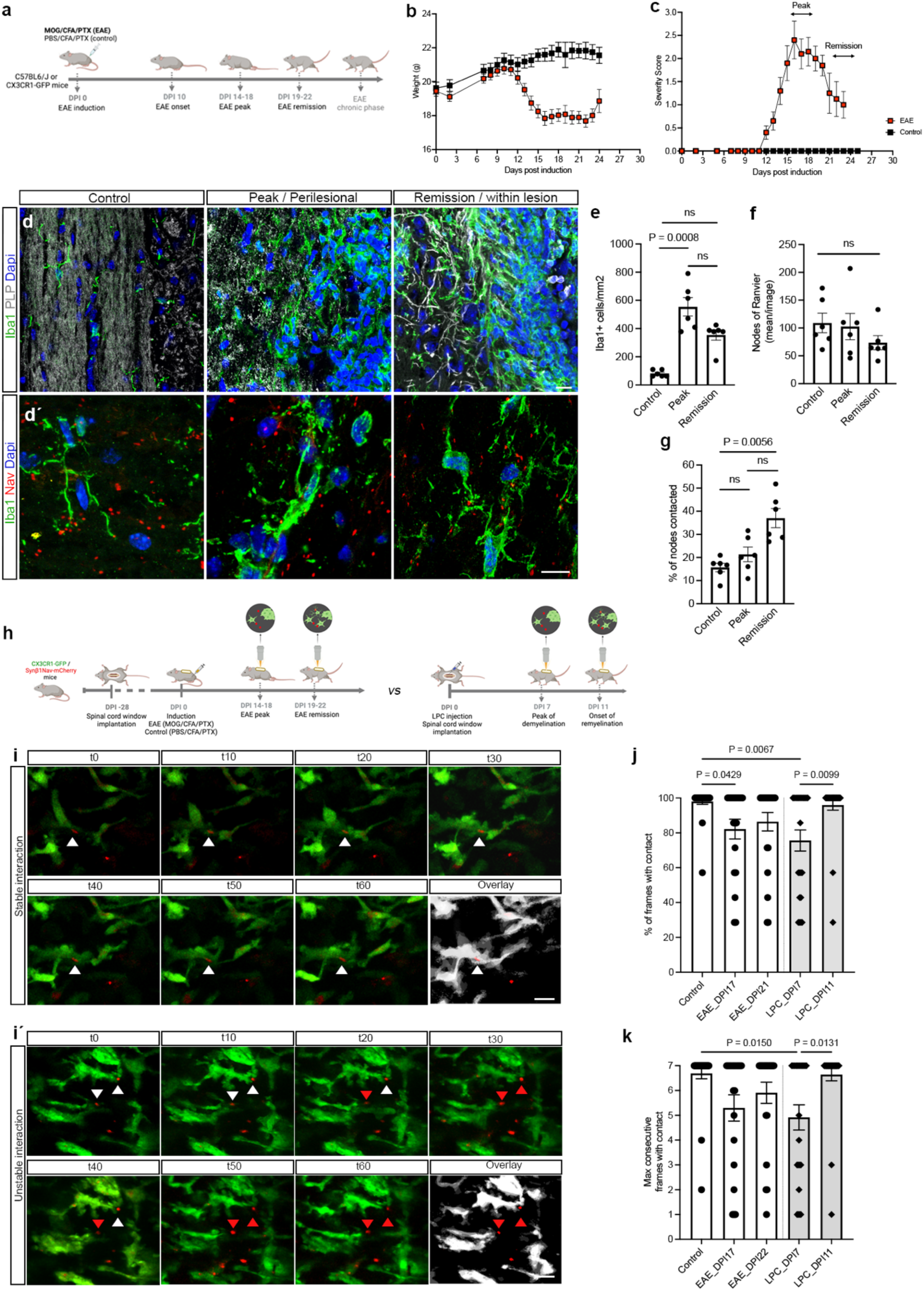
Remission is associated with an increased but heterogeneous microglia-node interaction in an inflammatory context. **(a)** Timeline of disease induction and follow-up in MOG₃₅₋₅₅ EAE mice and controls. **(b)** Body weight and **(c)** clinical score monitoring in EAE (n = 12) and control mice (n = 6) (mean ± SEM [standard error of the mean]). **(d)** Representative immunostainings of the dorsal spinal cord in control and EAE mice at the lesion border during peak disease and within lesions at the onset of remission. PLP staining (grey) for myelin and Iba1 (green) for microglia/macrophages (MDMs). Acute lesions at peak show high Iba1⁺ cell density with activated morphology (reduced ramifications and/or hypertrophied cell body), high DAPI density, and decreased PLP staining. Microglia–node interactions were analyzed in perilesional white matter (<250 µm). During remission, some lesions show partial resolution of inflammation, more ramified microglia, and partially myelinated axons at lesion edges. **(d΄)** Iba1⁺ cells (green) interacting with nodes of Ranvier (Nav, red) in the corresponding regions. **(e)** Iba1⁺ cell density/mm² across conditions (control: 81.4 ± 9.5 Iba1^+^ cells/mm^2^; peak: 554.5 ± 65.7; remission: 355.1 ± 37 in remission; n = 6 per condition). (f) Node density per analyzed area (mean ± SEM). **(g)** Percentage of nodes contacted across conditions (mean ± SEM); each dot represents the mean of 3–5 regions per animal. Quantification shows that 15.7 ± 1.8% of nodes are contacted in controls, 21.4 ± 3.1% at peak, and 37.1 ± 4.1% at remission onset; n = 6 animals per condition. **(h)** Schematic representation of the experimental design for *in vivo* live imaging in EAE and LPC-induced CX3CR1-GFP/Synβ1-Nav-mCherry mice. **(i-i΄)** Time-lapse imaging of the dorsal spinal cord (60 min movies; 1 frame/10 min) showing GFP⁺ microglia/MDMs (green) and nodes of Ranvier (mCherry, red), with **(i)** stable microglia–node interaction in control mice and **(i΄)** examples of unstable interaction in EAE mice. White arrowheads indicate a maintained microglia-node interaction while red arrowheads indicated a disrupted contact. **(j)** Percentage of frames per movie in which contact is maintained and **(k)** maximum number of consecutive frames showing maintained contact per movie; each dot represents one microglia/MDM–node pair in contact at t_0_ and tracked over 60 min. Control: 28 microglia-node pairs; EAE: 20 pairs (DPI17) and 22 pairs (DPI21); LPC: 24 pairs (DPI7) and 28 pairs (DPI11), from n = 6-9 mice. Statistical analysis: Kruskal-Wallis test with Dunn’s multiple comparisons in (e-g, j-k); ns, not significant. Scale bars: (d-d**΄**) 10 μm, (i-i**΄**) 5 μm.

In CX3CR1-GFP EAE mice, we combined stainings for GFP (resident and recruited microglia/macrophages) and CCR2 (infiltrating monocytes/macrophages, absent from microglia) (Fig. S1a), identifying GFP^+^/CCR2^-^ microglia, GFP^-^/CCR2^+^ monocytes, and GFP^+^/CCR2^+^ monocyte-derived macrophages (MDMs). At disease peak, MDMs and monocytes heavily infiltrated perilesional white matter (Fig. S1b). Whereas all nodal contacts were mediated by microglial cells in controls, there was a significant number of contacts mediated by MDMs in the perilesional tissue at disease peak (49,8% of nodes contacted), with fewer by monocytes (7.8%), and occasional dual contacts (Fig. S1c).

By contrast, at remission onset (48 h after peak, defined by EAE score decline), most nodal contacts were mediated by GFP^+^/CCR2^-^ cells (89.5% vs 8.5% by CCR2^+^ cells), with rare dual contacts (Fig. S2c). Combining staining for Iba1 with the microglia-specific marker TMEM119 (Fig. S2d), we found that at remission onset 97.4% of nodal contacts were made by TMEM119^+^/Iba1^+^ resident microglia (Fig. S2e–f), despite TMEM119^-^/Iba1^+^ cells (recruited myeloid cells and some activated microglia subsets) comprising 25.9% of Iba1^+^ cells (Fig. S2e-f) at this time point.

Together, these results suggest a predominant microglial involvement at nodes in homeostatic and repair contexts, whereas macrophage- and monocyte-mediated contacts are mostly confined to the acute inflammatory phase.

### Nodal contacts mediated by microglial cells are increased - but heterogeneous - at the onset of remission under inflammatory conditions

We previously reported that microglia-node contact frequency significantly increased at the onset of remyelination in a toxic demyelination model (focal spinal cord demyelination induced by lysolecithin) (37). To assess whether inflammation can affect neuron-microglia interactions at the nodes of Ranvier, we next investigated microglial contacts at the nodes in spinal cord white matter of EAE and control mice (Fig. 1a–d). We quantified this interaction as the percentage of nodes contacted at the peak of the disease, in the perilesional white matter adjacent to the acute lesions, and during remission (Fig. 1d-d΄).

At peak, we observed a 6.8 fold increase of microglia/macrophage cell pool in the perilesional tissue at peak compared to the control condition, followed by a trend towards reduction during remission (Fig. 1e). By contrast, nodal (Nav^+^, Fig. 1f) and paranodal (Caspr^+^, Fig. S2a-a΄) densities did not significantly vary between conditions in the studied area.

During remission, the percentage of nodes contacted was significantly increased, showing a 2.4-fold rise relative to control and a 1.7-fold rise relative to peak (Fig. 1g, Fig. S2b), independently of microglial cell and nodal densities (Fig. S2c-d). Although the magnitude of this increase observed at remission was slightly lower under inflammatory conditions compared to the toxic demyelination model (2.4-fold increase in EAE vs 3-fold in LPC (37)), microglia-node contact frequency in EAE exhibited a markedly greater interindividual variability.

### Microglia-node contacts remain unstable during EAE remission

To examine microglia/macrophage-nodal contact dynamics under inflammatory conditions, we next performed in vivo longitudinal time-lapse imaging in CX3CR1-GFP/Synβ1-Nav-mCherry mice. Contacts between GFP⁺ cells and axons at the node of Ranvier were analyzed in perilesional white matter surrounding acute lesions and in remyelinating areas in the dorsal funiculus of the spinal cord, during both EAE and LPC-induced demyelination. EAE mice were imaged at both peak and remission and compared with control (PBS/CFA/PTX-injected) and LPC-injected mice (Fig. 1h-i). In both EAE and LPC lesions, GFP⁺ cell-node interactions were destabilized in perilesional regions at peak (Fig. 1j-k). Interaction stability fully recovered in remyelinating LPC lesions (one-hour movies; 96% ± 3% (mean ± SEM) of timepoints with contact at the onset of remyelination vs. 76% ± 6% at demyelination peak). In contrast, no recovery was observed in EAE (86%±5% of timepoints with contact at the onset of remission vs. 82% ± 6% at EAE peak) (Fig. 1j-k). Further analysis in EAE mice showed that at peak, 45% of the existing contacts were lost during filming, compared with 32% during remission (Fig. S2e), with a mean detachment time of 30 minutes (Fig. S2f). Morphological analysis indicated that unstable contacts were more frequent among amoeboid GFP^+^ cells at peak (Fig. S2g-g΄΄), whereas morphology alone did not correlate with stability during remission (Fig. S2h-h΄). These results demonstrate that the stability of microglia-node interaction is durably impaired in an inflammatory context.

### High microglia-node interaction correlates with improved functional outcome during remission

At the onset of remission in EAE, we observed an important heterogeneity between animals in microglia-node interaction frequency (Fig. 1g). Since motor recovery from peak phase is also heterogeneous between animals, we next examined whether microglia-node interaction correlates with mice clinical outcome. To address this question, we conducted a multimodal analysis combining behavioral testing, clinical assessment and tissue analysis at the onset of EAE remission. To ensure the homogeneity of the group of animals studied, only mice reaching a severity score of 3 to 4 at peak were included in this study (“Severe EAE” group; 20 mice, from 5 separate EAE experiments) (Fig. 2a). Quantification of microglia-node interaction frequency in this cohort confirmed our initial observation of increased, but heterogeneous microglia-node interaction during remission (Fig. 2b-c), with between-subjects variance exceeding within-subjects variance. Furthermore, increased interaction correlated with lower disease severity score during remission for the EAE mice (Fig. 2d), improved recovery from relapse (as assessed by score reduction between peak and onset of remission, Fig. 2e) and greater grip-strength recovery from peak (Fig. 2f). To further investigate interindividual variability in microglia-node contact at remission and its impact on repair, we classified EAE mice according to microglia-node interaction as “high” or “low” contacting, based on a median split of the Severe cohort during remission (median value = 39.1% of nodes contacted, Fig. 2g-h). No significant differences in Iba1^+^ cell density (Fig. 2i) or nodal density (Fig. 2j) were observed between the “low” and “high” contacting animals. High-contacting animals were confirmed to exhibit significantly lower severity scores at the last follow-up compared with low-contacting mice (Fig. 2k). Taken together, these observations reveal a strong interindividual heterogeneity in microglia-node interaction at the onset of remission, increased interaction being associated with improved recovery.

**Figure 2.**
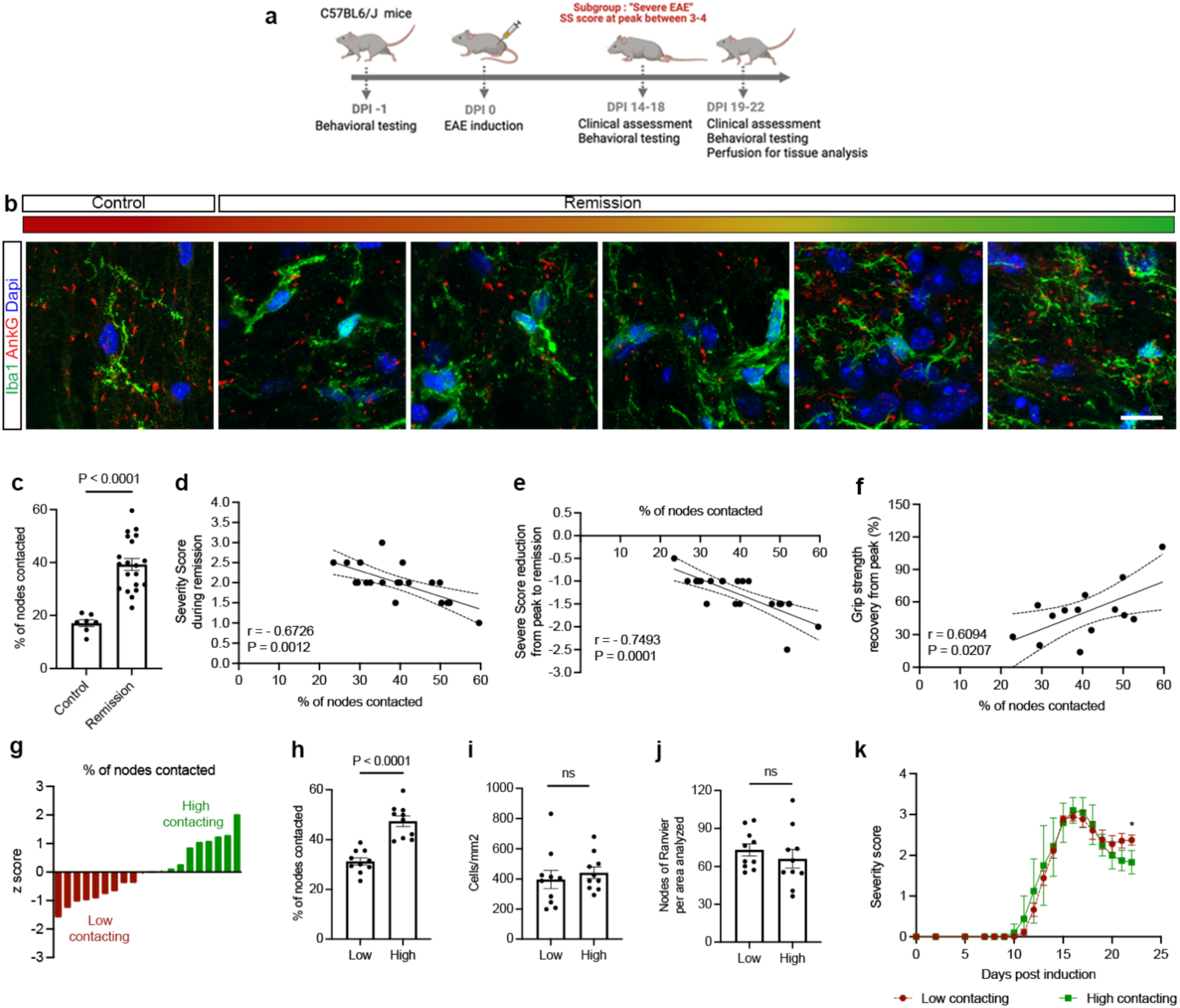
Increased microglia-node interaction during remission is associated with improved functional recovery in severe EAE. **(a)** Following EAE induction in C57BL/6J female mice, animals reaching a severity score ≥3 at peak were classified as “Severe EAE” and followed until remission. **(b)** Examples of microglia/MDM-node interactions in control spinal cord white matter tracks and in sub-acute lesions at remission, ranging from lower frequency (∼25%, red) to higher frequency (∼75%, green). **(c)** Frequency of microglia/MDM–node contacts in control vs EAE remission (mean ± SEM, n = 20 EAE mice, n = 7 controls; each dot represents the mean of 3–5 analyzed areas per animal). **(d–f)** In the Severe EAE group, increased microglia–node interaction correlated with improved outcomes: **(d)** reduced severity score during remission (n = 20), **(e)** increased recovery (change in severity score from peak to remission, n = 20), and **(f)** improved grip-strength recovery from peak (n = 14). **(g)** Median split of the cohort during remission (median = 39.12%) separated mice into “low” (n = 10) vs “high” (n = 10) contacting groups (shown as z-scores). **(h)** Percentage of nodes contacted per condition (mean ± SEM) in low- vs high-contacting groups. **(i)** Iba1⁺ cell density/mm² (mean ± SEM). **(j)** Node of Ranvier density per analyzed area (mean ± SEM). **(k)** Severity score at last follow-up was significantly higher in low-contacting mice. Statistical tests: t-test with Welch’s correction when applied in (c, h-i), Spearman correlations in (d–e), Pearson correlation in (f), Solid lines represent linear regression fits with dashed lines showing 95% confidence intervals. Two-way ANOVA in (k). *: p < 0.05; ns: not significant. Scale bars: 20 μm (b).

### Microglia-node interaction varies depending on local immune cues

To characterize molecular cues that could impact microglia-node interaction and contribute to its heterogeneity during remission, we next performed Luminex-based multiplex profiling of the cytokines and chemokines present in spinal cord tissue from Severe EAE mice (n = 12) (Fig. 3a). Unsupervised clustering identified three major inflammatory profiles (Groups A–C) (Fig. S3a). Group A displayed a mixed Th1/Th17–Th2 signature (GM-CSF, granzyme B, leptin, IL-6R, IL-4, IL-13, IL-9, IL-19, IL-10, IL-31, IL-27, CCL25 [TECK], M-CSF) together with ENA-78, VEGFR2 and RANKL expression, suggesting tissue remodeling (Fig. S3b). Group B presented globally lower cytokine and chemokine expression compared to Group A and C, while group C showed reinforced expression of molecules associated with peripheral immune cell recruitment, such as IP-10/CXCL10, CCL5, CCL2 and CCL7, compared to Group A (Fig. S3c). Importantly, mice in Group A exhibited significantly higher microglia-node contact frequency than mice in Groups B and C (Fig. S3d).

**Figure 3.**
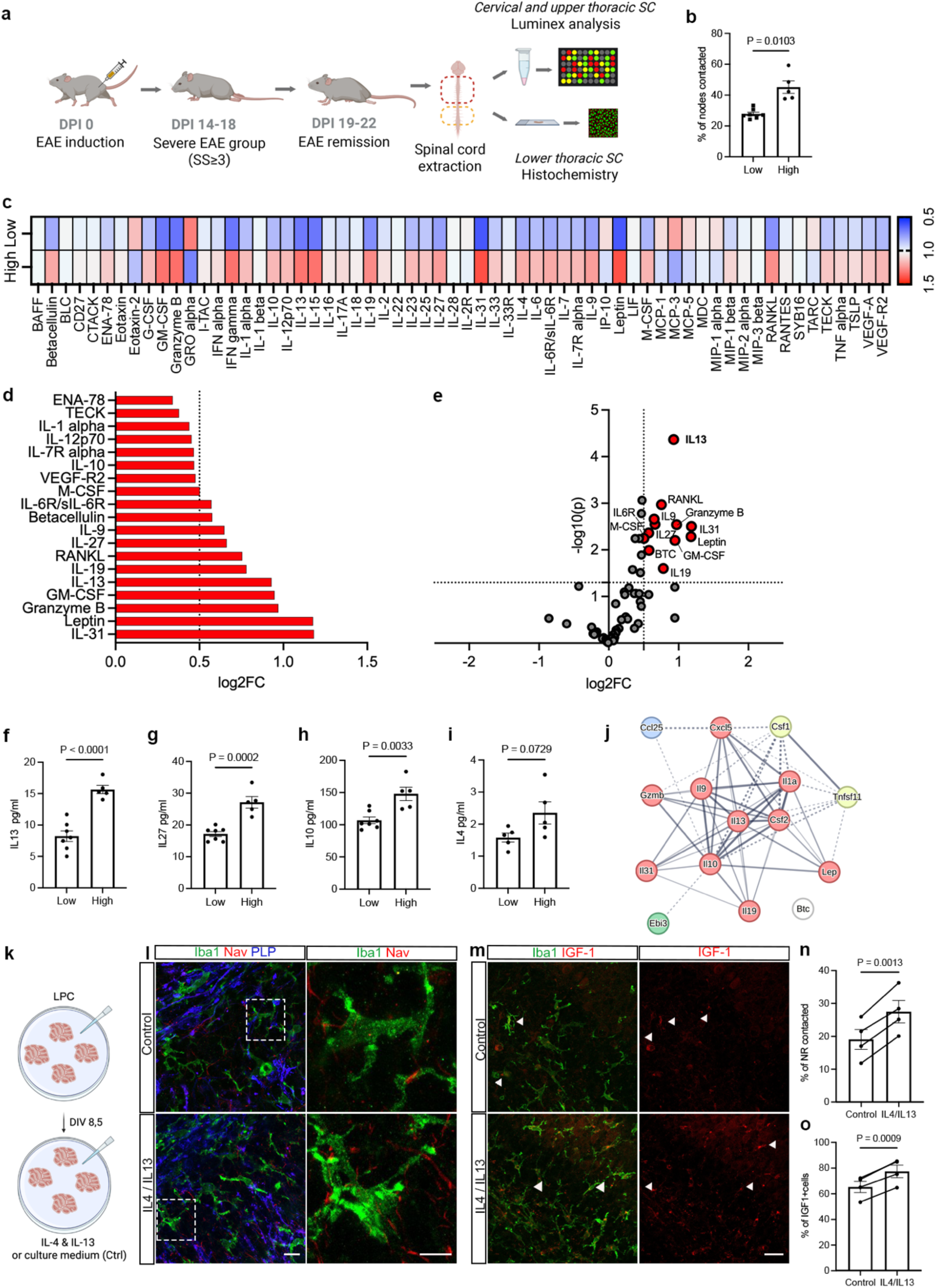
Th2/Treg responses are associated with increased interaction during remission. **(a)** Following EAE induction in C57BL/6J female mice, animals that reached a peak clinical score of ≥3 were classified as having Severe EAE and monitored through remission. Brain and spinal cord tissues were collected from each mouse. The brain together with the cervical and upper thoracic spinal cord segments were processed for Luminex analysis, whereas the lower thoracic to lumbar spinal cord segments were used for immunohistochemistry to validate microglia–node interactions in each individual animal. **(b)** Frequency of microglia-node of Ranvier contacts in low- vs high-contacting mice (n = 5-7 mice/group; each dot = mean of 3-5 areas analyzed per mouse; bars = mean ± SEM). **(c)** Heatmap of differentially expressed cytokines and chemokines in the spinal cord of low- (n = 5) vs high-contacting mice (n = 7). Values are log₂ fold change (log2FC) relative to low-contacting mice. **(d)** Enriched expression of molecules in high-contacting compared to low-contacting animals. **(e)** Volcano plot of cytokines and chemokines differentially expressed in high-contacting mice at remission. Red dots highlight significant upregulation (p < 0.05, log₂FC > 0.5). **(f-i)** Quantification of expression of **(f)** IL-13, **(g)** IL-27, **(h)** IL-10 and **(i)** IL-4 in low- vs high-contacting mice (mean ± SEM). **(j)** Protein–protein interaction (PPI) network of upregulated cytokines. A PPI network was generated using STRING and visualized in Cytoscape (automatic k-mean clustering, colors indicate different clusters, red=JAK-STAT signaling pathway (IL-13 in the center of the cluster, Csf2, IL-10, IL-9, IL-19, IL-1a, Leptine, Gzmb:Granzyme B, IL-31, Cxcl5), blue=chemotactic activity on macrophages and dendritic cells (CCL25), green=T lymphocyte regulatory signaling (Ebi3 : IL-27), yellow = T cell proliferation and differentiation pathway (Csf1, Tnfsf11). **(k)** Schematic representation of *ex vivo* model of treated demyelinated organotypic cerebellar slices. Briefly, 6-8 sagittal cerebellar slices from post-natal day 9 (P9) old C57BL6/J mice were plated in two membranes and put in culture. Slices were demyelinated with LPC at day *in vitro* (DIV) 6, for 15 h 30 min. At DIV 8.5, half the slices from one animal (one membrane) were incubated with 20 ng/mL IL-4 and 20 ng/mL IL-13 for 24 h, followed by fixation and immunostainings at DIV 9.5, while the other half were used as control. **(l)** Immunostainings of Iba1^+^ cells (green), nodal axonal domains (Nav, red), and myelin (PLP, blue) in treated and untreated demyelinated slices. **(m)** Immunostainings of Iba1^+^ cells (green) and IGF-1 (red) in treated and untreated demyelinated slices. **(n)** Quantification of the percentage of nodes of Ranvier (NR) contacted in control and treated slices (n = 4 animals/condition; paired slices per animal; mean ± SEM). **(o)** Quantification of distribution of IGF-1^+^ microglial cells in control and treated slices (n = 4 animals/condition; mean ± SEM). Statistical analysis: t-test (with Welch’s correction when applicable) in **(b, g-j)**, paired t-test in **(n-o)**. Scale bars: 25 μm **(l-m)**.

We then performed a differential expression analysis comparing high- vs low-contacting animals (as defined above) to assess whether they associate with specific cytokine and chemokine profiles. Among the 64 analytes quantified, 19 were statistically significantly differentially expressed in high-contacting (n = 5) vs low-contacting (n = 7) mice (p < 0,05; Fig. 3 c), including GM-CSF, IL-1α, IL-12, IL-6R, IL-7Rα, granzyme B, IL-9, IL-19, IL-13, IL-10, IL-27, IL-31, RANKL, betacellulin, VEGF-R2, leptin, M-CSF, CCL25 and ENA-78. Of these, 12 were significantly overexpressed in the high-contacting group (log2FC > 0,5; Fig. 3d-e): 4 rather associated with a Th1/Th17 profile (GM-CSF, granzyme B, leptin, IL-6R, Fig. S3e-g) and 8 involved in Th2 and/or Treg responses (IL-13, IL-9, IL-19, IL-27, IL-31, M-CSF, RANKL, betacellulin, Fig 3f-h and FigS3h-j), with IL-4 showing a trend towards upregulation (log2FC = 0,6, p = 0,0729; Fig 3i). Overall, the immune secretome of high-contacting mice reflected a mixed adaptive immune signature, with a shift towards Th2-skewed and regulatory/myeloid-modulating responses.

Gene ontology enrichment analysis of the 19 upregulated molecules in high-contacting mice revealed strong activation of the JAK-STAT signaling pathway (‘regulation of receptor signaling pathway via JAK-STAT’, ‘positive regulation of receptor signaling pathway via JAK-STAT’, ‘positive regulation of tyrosine phosphorylation of STAT protein’; GO Biological Process 2025, FDR < 0.05; Fig. S3j). Protein-protein interaction (interactome) analysis of the upregulated molecules in high-contacting mice revealed a highly interconnected network centered on IL-13 and CSF2, with intricate connections to IL-10, IL-9 and IL-19 (Fig. 3j), supporting IL-13-driven STAT6 activation, known to promote IGF-1-expressing, pro-regenerative microglial states.

IL-13 emerged as a potential key factor as it (i) was the most robustly overexpressed cue in high- vs low-contacting animals (Fig. 3i), (ii) localized at the center of the highly connected network of upregulated molecules in high-contacting animals (Fig. 3j) and (iii) is further known to modulate neuronal physiology (40–43). Therefore, we next investigated whether its presence could promote microglia-neuron interaction at nodes. IL13 is known to act synergistically with IL4 (44), which also showed a tendency towards increased expression in the high-contacting EAE mice. We therefore added exogenous IL-13/IL-4 at the peak of LPC-induced demyelination in organotypic cerebellar slices, an accessible model previously validated for microglia-node interaction study (Fig.3k, (37, 38)). A 24h IL-13/IL-4 treatment reinforced microglia-node interaction by 44% compared to control condition (Fig. 3l, n) and was further associated with a 19% increase in IGF-1^+^ expressing microglia compared to control condition (Fig. 3m, o). Altogether, these data indicate that microglia-node contacts are modulated by cues secreted by adaptative immune cells during lesion resolution, and this modulation is further linked to specific microglial signatures.

### Tissue repair is increased in high microglia-node interaction context

We next examined *in vivo* whether increased microglia-node contact frequency during remission was also associated with phenotypic changes of microglial cells. We first performed iNOS/Iba1 and IGF1/Iba1 stainings. While the percentage of the pro-inflammatory iNOS⁺/Iba1⁺ cells did not significantly differ in regard to contacts (Fig. 4a-a΄), the percentage of the pro-regenerative IGF-1⁺/Iba1⁺ cells was increased by 1.6-fold in high vs low contacting mice (Fig. 4b-b΄). We next performed triple labelling for Iba1, iNOS and IGF-1 (Fig. S4a), which showed that 32.3%± 4.3% (mean ± SEM) of Iba1^+^ cells co-expressed iNOS and IGF-1, while 58,1%± 4.3% were positive for iNOS only and 2,7% ± 1.3% for IGF-1 only, indicating that most IGF-1⁺ cells were in a transitional state co-expressing iNOS at the stage of EAE considered (Fig. S4b). Only a small fraction of Iba1^+^ cells were negative for both iNOS and IGF1 (Fig. S4b). Looking at the repartition of activated microglia expressing iNOS and/or IGF1 in high- vs low-contacting animals, we observed a significantly decreased iNOS^+^-only population, associated to a significantly higher proportion of iNOS⁺/IGF-1⁺ double-positive cells in high-contacting animals, supporting a shift towards pro-regenerative microglia in these mice (Fig. 4b΄΄).

**Figure 4.**
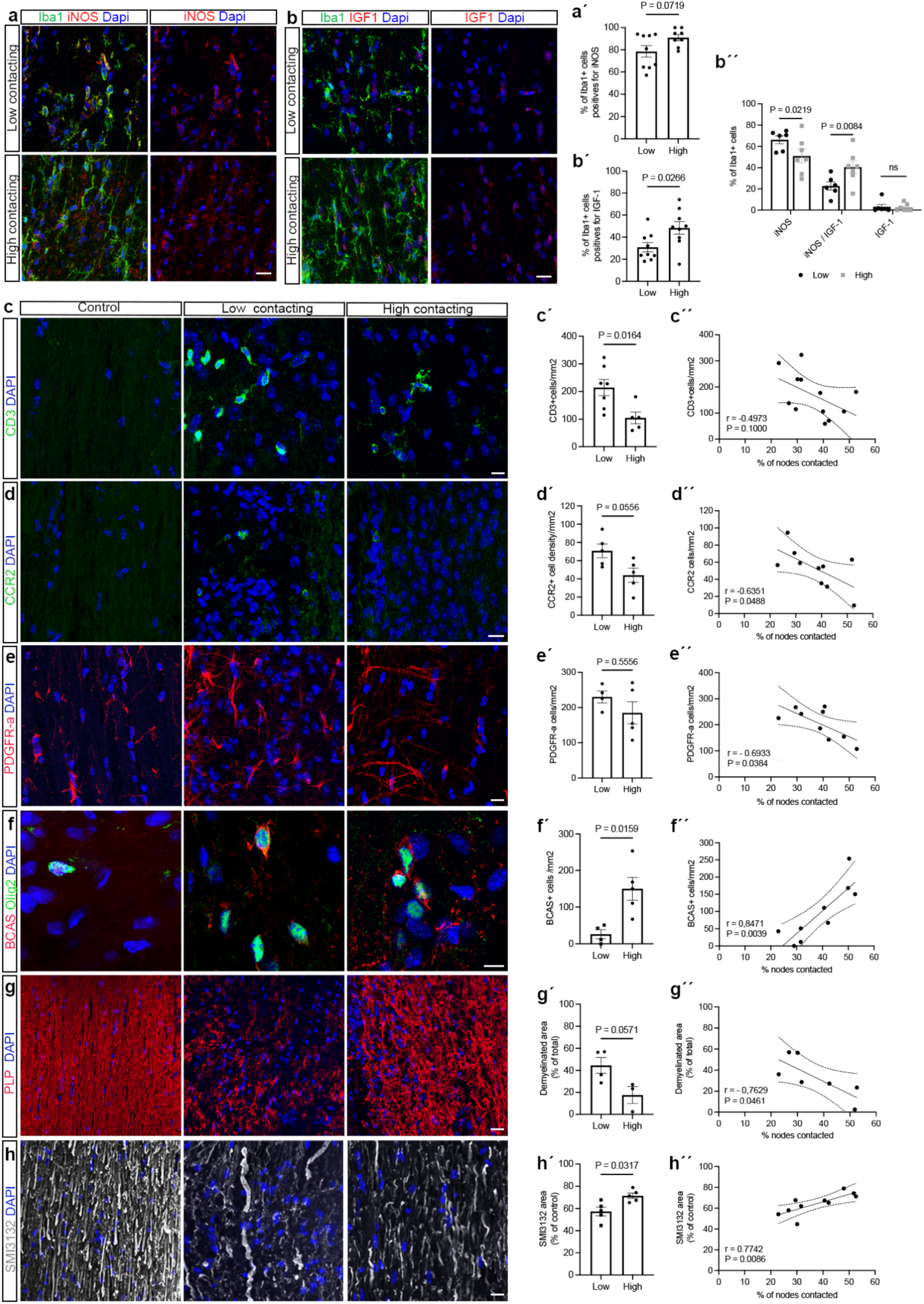
Reinforced microglia-node interaction during remission correlates with better tissue repair. **(a)** Representative images of iNOS^+^ cells (red) and **(b)** IGF-1^+^ cells (red) amongst Iba1^+^ cells (green) during EAE remission in low- vs high- contacting mice spinal cord. Frequency of **(a΄)** iNOS^+^ and **(b΄)** IGF-1^+^ cells among all Iba1^+^ cells in the two groups. The percentage of IGF-1^+^ cells was increased in the high-contacting group (48.3% ± 5.7 vs 30.9 ± 4.2, n = 9/group). **(b΄΄)** Frequency distribution of iNOS^+^, IGF-1^+^, and iNOS^+^/IGF-1^+^ cells amongst Iba1^+^ cells in high- and low-contacting animals. Representative immunostainings of: **(c)** CD3^+^ cells (green), **(d)** CCR2^+^ cells (green), **(e)** PDGFRα^+^ cells (red), **(f)** BCAS^+^ (red) and Olig2^+^ cells (green), **(g)** myelin staining (PLP, red), and **(h)** axonal covered area (SMI31 and SMI32, grey) in control and in the two EAE subgroups. **(c΄-h΄)** Density distributions between low- and high-contacting animals for: **(c΄)** CD3^+^ cells (n = 5-7/group), **(d΄)** CCR2^+^ cells (n = 5/group), **(e΄)** PDGFR-α^+^ cells/mm^2^ (n = 4-5/group), **(f΄)** BCAS^+^ cell/mm^2^ (n = 4-5/group), **(g΄)** demyelinated area (% of total area, n = 3-4/group), and **(h΄)** axonal density distribution (n = 5/group). **(c΄΄-h΄΄)** Pearson correlations between cell densities and the frequency of microglia–node interaction during remission; solid lines show linear regression fits and dashed lines indicate 95% confidence intervals. Axonal and myelin density are expressed as % of SMI31/32 or PLP covered area relative to the mean covered area of n = 3 control mice. Statistical analysis: Mann-Whitney test in **(a΄, c΄-h΄)**, Welch’s t-test in **(a΄΄, b΄)**, two-way ANOVA in **(b΄΄)**, Pearson correlations in **(c΄΄-h΄΄)**. ns, not significant. Scale bars: 25 μm in **(a-e)**, 5 μm in **(f)**, 50 μm in **(g)**.

We further quantified CD3⁺ T cell retention in the spinal cord at remission and found that high-contacting mice displayed a T cell density reduced to half compared to low-contacting mice (Fig. 4c-c΄΄). Similarly, CCR2⁺ monocyte/macrophage densities inversely correlated with microglia-node contact (Fig. 4d-d΄΄ and statistical table), indicating that high microglia–node interaction is associated with a microenvironment characterized by the reduced presence of peripheral immune cell.

Given that IGF-1 expressing microglia have been directly implicated in oligodendroglial lineage modulation (18), we next examined OPC and newly-formed pre-myelinating oligodendrocyte distribution. OPC density presented no significant difference between low- and high-contacting animals, though a moderate negative correlation between microglia–node interaction rate and PDGFRα⁺ OPC density was observed (Fig. 4e-e΄΄). We further assessed the generation of new oligodendrocytes in high vs low contacting EAE mice, by staining for BCAS, which is described as a marker of newly differentiated oligodendrocytes (45). BCAS⁺/Olig2⁺ newly differentiated oligodendrocytes–presented a significant 5.7-fold increase in high-contacting compared to low-contacting condition and positively correlated with microglia–node interaction (Fig. 4f-f΄΄). Consistent with enhanced oligodendrogenesis, demyelinated lesion area was decreased by 60.3% in remission (Fig. 4g-g΄΄), while axon-covered area significantly increased by 24.6% in high vs low contacting animals.

To assess the influence of multiple variables on microglia-node interaction, a partial least squares regression (PLSR) analysis was performed for descriptive purposes using clinical and biological variables from Severe EAE group. Microglia-node interaction frequency best discriminated the cohort into two groups (Fig. S4c). High interaction levels were associated with increased IGF-1⁺ cell density, axonal density, and clinical recovery (both clinical score improvement and grip strength recovery), whereas high OPC density, demyelinated area, and severity score at remission correlated with low interaction (Fig. S4d). The presence of iNOS⁺ microglia did not show a notable association with microglia-node interaction.

Altogether, high-contacting mice exhibited a less inflammatory lesion environment, enhanced repair and improved clinical outcome, including lower severity scores and better strength recovery from peak disease.

### Low-intensity exercise post-peak reinforces microglia-node interaction and improves recovery

There is growing evidence that exercise and motor learning positively influence myelination (46–50). Exercise prior to the induction or onset of EAE has further been shown to decrease inflammatory cytokine secretion, reduce demyelination, and promote IGF1 secretion (51–55). We also recently showed that a physiological increase in neuronal activity can promote microglia-node interaction at the onset of repair, with a beneficial outcome on microglial signature and repair (38). Therefore, we wondered whether physical exercise, when initiated after the peak of EAE, could modulate microglia-node interaction and promote repair. We induced EAE in mice previously habituated to slow walking on a treadmill and mice reaching a severity score of 2 or 2.5 at peak were randomly assigned to either the physical activity or the sedentary group. The physical activity group performed low-intensity exercise (LIE) from 48h post-peak for 4 consecutive days, starting post-peak, while mice of the sedentary group (SED) were placed on a non-running treadmill for similar timings and duration (Fig. 5a). The treadmill speed was chosen to match a walking pace for a mouse with paraparesis, as determined by free locomotion measurements on a separate anterior open-field test. Severity score follow-up in the LIE vs SED groups showed improved recovery in the LIE group from the second day of exercise (Fig. 5b). Automated gate recordings were performed at baseline prior to EAE induction, and during the first and last exercise sessions. Multiple gate parameters are altered during EAE, with swing time reflecting subtle changes in myelination (50) and being modified at EAE onset (56, 57) (Fig. 5c). In our experiments, swing time increased by ∼45% at the peak of EAE in both groups. Physical activity led to a significant 25% reduction in swing time, restoring performance to near-baseline levels, whereas sedentary mice showed only a smaller, non-significant decrease (Fig. 5d-d΄). Overall, the recovery in swing time from first to last session was significantly more pronounced in the LIE group (Fig. 5d΄΄).

**Figure 5.**
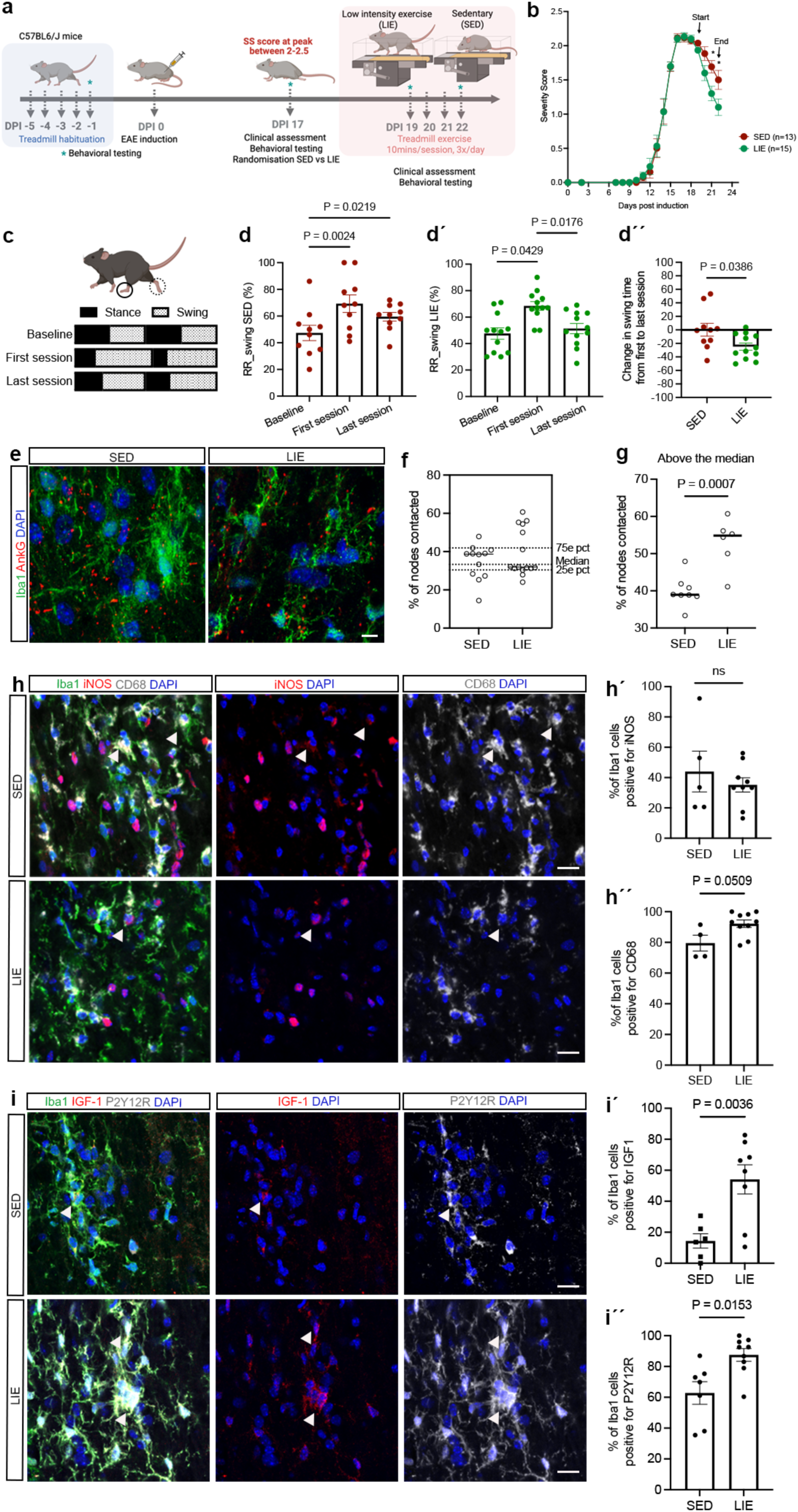
Low intensity exercise following relapse reinforces microglia/macrophage interaction with the node of Ranvier and improves outcome. **(a)** Schematic representation of the experimental protocol. Following EAE induction, mice were monitored until peak; those reaching a severity score of 2–2.5 were randomly assigned to sedentary (SED) or low-intensity exercise (LIE) groups and completed the 4-day protocol. Automated gait recordings were performed during the first and last exercise sessions. **(b)** Severity score over time in SED and LIE groups (mean ± SEM, n = 28 animals from 4 independent EAE experiments). **(c)** Schematic representation of the percentage of time spent in stance and swing phases during baseline, first, and last sessions. **(d)** Swing time of the right rear paw (RR) for SED mice (red) and **(d΄)** LIE (green) mice at baseline, first, and last sessions (mean ± SEM, n = 10 mice/condition). **(c΄΄)** Change in swing time (first to last session) for each animal (mean ± SEM, n = 10 mice/condition). **(e)** Representative images of microglia/macrophage–node interactions (Iba1, green; AnkG, red). **(f)** Frequency of microglia/macrophage–node interactions in SED vs LIE (each dot = mean of 5 areas/mouse; bars = mean ± SEM). Median and interquartile range (25^th^-75^th^ percentiles) are indicated on the right y-axis. **(g)** Due to differences in distribution between groups, with a subset of LIE mice showing reinforced interactions, a median split was applied. Mean frequency of contacts was compared between SED and LIE mice above the median. **(h)** Representative immunostainings of iNOS and CD68 Iba1^+^ cells. **(h**΄) Proportion of Iba1⁺ cells expressing iNOS (mean ± SEM, n = 5-10 mice/condition), and **(h΄΄)** Proportion of Iba1⁺ cells expressing CD68 (mean ± SEM, n = 4-10 mice/condition). **(i)** Representative immunostainings of IGF-1 and P2Y12R Iba1^+^ cells. **(i**΄) Proportion of Iba1⁺ cells expressing IGF-1 (mean ± SEM, n = 6-8 mice/condition) or **(i**΄΄) P2Y12R (mean ± SEM, n = 7-9 mice/condition). Statistical analysis: Kruskal-Wallis test with Dunn’s multiple comparisons in **(c-c΄)**, Mann–Whitney test in **(g, h**΄΄), Welch’s t-test in **(d, g**΄**, h΄, i΄, i΄΄)**. ns, not significant. Scale bars: **(e)** 10 μm; **(h, k)** 50 μm.

Microglia-node contact study revealed only a tendency towards increase of the mean contact frequency between the SED and LIE groups at the timepoint considered. However, a third of LIE mice showed markedly increased interaction (mean microglia-node interaction frequency > 50%), whereas no SED mice displayed this level of interaction (p = 0.0267, statistical table). Furthermore, the mean percentage of nodes contacted was significantly higher in both high- and low-contacting cohorts in LIE compared to SED mice (i.e. above- and below-median groups respectively, Fig. 5f-g and statistical table), suggesting that LIE led to increased contacts for both low- and high-contacting populations. There were no significant differences in the expression of pro-inflammatory markers such as iNOS or CD68 (Fig. 5h–h΄) between LIE and SED animals, but the LIE group showed a significant increase in IGF-1^+^ innate immune cells, as well as in Iba1⁺ cells expressing the homeostatic marker P2Y12R (Fig. 5i–i΄΄).

These results support the idea that physical exercise can lead to a shift towards enhanced microglia-node interaction, potentially promoting a switch towards pro-regenerative microglial signatures and accelerating their return towards homeostatic states.

### Microglia-node of Ranvier interaction is reinforced in shadow plaques of MS patients and correlates with remyelination

As microglia interact with the node of Ranvier in the human brain (37), we next investigated whether this interaction is modulated in patient with MS (pwMS). To address this, we analyzed post-mortem hemispheric white matter tissue from six pwMS and four healthy donors (Supplementary table 1) and classified lesions according to the updated histological classification (58) (Fig. S5a). We examined how resident microglial cells (TMEM119^+^) interact with the node of Ranvier in control tissue, as well as in normal-appearing white matter (NAWM) and remyelinating shadow plaques from pwMS patients (Fig. 6a–c). Interaction frequency was similar in NAWM and healthy donors, but significantly increased in shadow plaques (by 78,4% compared to NAWM and by 70% compared to control subjects, Fig. 6d). Consistent results were obtained for homeostatic P2Y12R⁺ microglia (Fig. 6e–f), and this effect was further independent of both nodal and microglial cell density (Fig. S5b–d).

**Figure 6.**
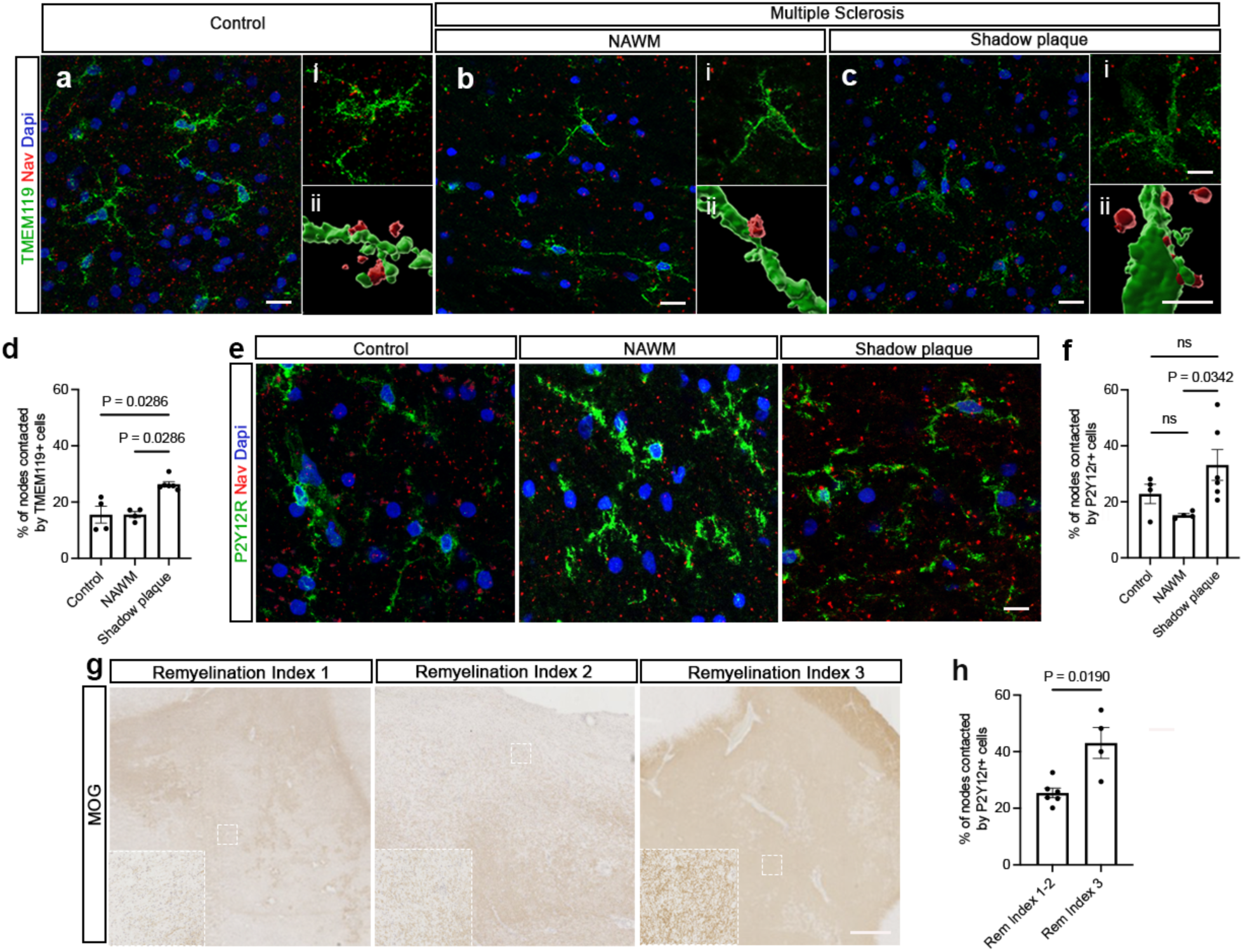
Microglia-node of Ranvier interaction is reinforced in shadow plaques of MS patients. **(a–c)** Fluorescent immunostainings in human post-mortem hemispheric white matter from control tissue **(aᵢ, bᵢ)**, MS normal-appearing white matter (NAWM) **(aᵢᵢ, bᵢᵢ)**, and shadow plaques **(aᵢᵢᵢ, bᵢᵢᵢ)**. Representative lower-magnification images show resident microglia (TMEM119⁺, green) interacting with nodal domains (Nav, red); contact sites are indicated with asterisks. Insets: zoomed microglia–node contacts and corresponding 3D reconstructions. **(d)** Quantification of TMEM119⁺ cell–node contacts (26.4 ± 0.9% of nodes contacted in shadow plaques vs 14.8 ± 0.9% in NAWM and 15.5 ± 3.0% in control subjects; n = 4–6 lesions/condition). **(e)** Fluorescent immunostaining showing homeostatic microglia (P2Y12R⁺, green) in control, NAWM, and shadow plaques. **(f)** P2Y12R⁺ cell–node contacts, 33.0 ± 5.4% of nodes contacted in shadow plaques vs 15.2 ± 0.6% in NAWM and 22.8 ± 3.4% in control, n = 4–7 lesions/condition. **(g)** For remyelinating lesions, the extent of remyelination was determined semi-quantitatively (59), and lesions were classified using a remyelination index: 1 (10–50% MBP⁺ area), 2 (50–90% MBP⁺ area), and 3 (>90% MBP⁺ area). **(h)** Microglia–node interaction frequency is increased in lesions with remyelination index 3 compared to index 1–2 (25.40 ± 1.70% of nodes contacted in remyelination index 1-2 vs 43.10 ± 5.50% in remyelination index 3, p = 0.019; n = 4–6 lesions/condition). Statistical analysis: Kruskal-Wallis test with Dunn’s multiple comparisons correction in **(d, f)**, Mann-Whitney test in **(h)**; ns, not significant. Scale bars: 20 μm in **(a–c, e)** and 500 μm in **(g)**.

Because the extent of remyelination varies across lesions (59), shadow plaques were further classified according to the proportion of remyelinated area (remyelinating index [rem. index]; 1 = 10–50% remyelinated, 2 = 50–90%, 3 > 90%) (Fig. 6g). High levels of interaction between homeostatic microglia and nodes of Ranvier were 1.7-fold more prevalent in areas with high remyelination (rem. index 3) compared to areas with lower remyelination indices (Fig. 6h). In conclusion, microglial cells interact with nodes of Ranvier in both healthy and pwMS human brains, and these interactions are enhanced in remyelinating lesions. This supports the idea that microglia-node contacts may contribute to repair processes, consistent with findings from demyelinating mouse models.

## Discussion

Microglia-neuron crosstalk at nodes of Ranvier shapes microglial signatures in repair (38). Here, we show that inflammation tunes microglia-node interaction in EAE, as high contact frequency at remission onset correlates with a distinct cytokine profile, including elevated IL-13. High microglia-node interaction further associates with improved repair in both pwMS tissue and EAE, and low-intensity exercise during remission enhances this interaction, promoting pro-regenerative microglia and functional recovery. Together, these data support microglia-node interaction as a key component of a positive feedforward loop driving CNS repair.

### The nodes of Ranvier are dynamically engaged by distinct innate immune cells across EAE time course

The nodal area has previously been reported to be contacted by monocyte-derived macrophages (MDMs) at the onset of EAE, where these cells are thought to disrupt paranodal junctions and initiate demyelination, while activated microglia are engaged in other processes such as debris clearance (39, 60). At the peak of EAE, we observed MDMs/monocyte-mediated contact at nodes near acute lesions, which may contribute to the outward lesion propagation observed in EAE (61, 62). In contrast, at the onset of remission in EAE, microglial cells account for the vast majority of nodal contacts, suggesting a role in repair at this stage. These changing patterns support non-redundant roles of MDMs and microglia after demyelination (39, 63, 64). The local signals mediating macrophage contact at the node are still unknown, though potassium may play a role, as MDMs are known to express the 2-pore potassium channel THIK-1 (65), which contributes to microglial potassium read-out (37, 38, 66, 67). A small number of nodes are simultaneously contacted by microglia and monocyte/MDM, further supporting the concept of the node of Ranvier as a “communication hub” (37, 68), that may coordinate the resident-recruited immune crosstalk (31, 39, 69–71).

### The inflammatory context alters microglia-node interaction

Longitudinal spinal cord imaging in EAE and LPC models showed that EAE induces a persistent alteration of microglia-node contact stability, whereas it is restored during early remyelination after LPC. At the peak of EAE, unstable contacts coincide with amoeboid, highly activated microglia, whereas by remission, morphology no longer predicts contact behavior, implying distinct mechanisms of microglia-nodal contact dysregulation over time.

The chronic inflammatory milieu in EAE is a likely contributor to altered microglia-node interaction. Lymphocyte-derived cytokines and chemokines, including IFN-γ, TNF-α, IL-1β, IL-6, CCL-19, and GM-CSF are potent drivers of pro-inflammatory microglial states (31, 72–75). Inflammation is also known to impact neuronal integrity and induce early nodal pathology in MS and its models (60, 76–81). This, together with the pathological neuronal firing patterns observed in EAE (82), may further contribute to alter microglia-node interaction and its functional outcome. Abnormal firing itself can indeed maintain pro-inflammatory microglia after demyelination (38), suggesting a self-reinforcing loop where inflammation distorts neuronal activity, destabilizes microglia-node contacts and blocks the shift towards pro-regenerative microglia.

Interestingly, in EAE, both the stability and frequency of microglia-node interaction were highly heterogeneous across animals. Mice with high contact frequency displayed a distinct immune profile shifting from Th1 towards Th2 and, to a lesser extent, Treg-associated signatures, with upregulation of IL-13, IL-9, IL-19, IL-10 and IL-27 (83) which are associated to repair. In contrast, low-contact mice express chemokines that attract peripheral immune cells, with high levels of eotaxin-2 and MCP-1 (CCL7), associated to recruitment of eosinophils and neutrophils (84) and lymphocytes and monocytes (85) respectively. Both are known to aggravate EAE (86, 87) and high eotaxin-2 levels have been observed in the CSF of patients with inflammatory demyelinating disorders and significantly correlate with disability progression (88).

We identified IL-13, the most strongly upregulated cue in the spinal cord of high-contacting animals, as a potent regulator of microglia-neuron interaction at the nodes of Ranvier. Exogenous administration of IL-13, in combination with IL-4, which acts synergistically with IL-13 (44), was sufficient to enhance microglia-neuron interaction *ex vivo* following demyelination. IL-13 and IL-4 have been shown to enhance the excitability of sensory neurons in dorsal root ganglia (43) and IL-13 can promote phosphorylation of NMDA and AMPA receptors following CNS injury (41), suggesting a functional link between immune signaling and neuronal activity that may facilitate microglia-neuron interaction. IL-13/IL-4 treatment further led to an expansion of IGF-1^+^ microglia, consistent with IL-13 role in shaping microglial phenotype (18, 89). In line with this, we observed an enrichment in a transitional iNOS⁺/IGF1⁺ phenotype in high contacting animals. As IGF-1^+^ microglia are more likely to engage nodal domains than IGF-1^-^ activated microglia (37), this suggests IL-13 participates in a positive feedforward loop linking nodal interaction and microglial pro-regenerative polarization.

High-contact animals also overexpress IL-27, produced by microglia and astrocytes, as well as IL-10, which can promote neuroprotective, pro-regenerative microglial programs and restrains pathogenic T cell responses, including Th17, similar to IL-19 (90–99). In pwMS, IL-27 is enriched at active lesion borders (100), where remyelination attempts concentrate (59), further linking this cytokine to repair.

Altogether, these findings point to a coordinated immune program, that stabilizes microglia-node interactions and promotes lesion resolution, a concept that warrants deeper transcriptomic dissection in high- vs low-contact animals. The interindividual heterogeneity observed in EAE echoes the variability seen in multiple sclerosis patients, where the density and functional signatures of T lymphocytes are known to influence repair (101–103). Overall, high nodal microglia-neuron interaction defines a favorable repair context, whereas low interaction associates with poorer tissue repair and worse clinical evolution.

### Exercise post-peak reinforces microglia-node interaction and promotes repair

An emerging body of works suggests that motor learning and/or physical activity is beneficial in MS and its models (48, 104–106). Exercise performed prior to EAE induction, or during the acute symptomatic phase, has been shown to reduce immune cell infiltration, decrease secretion of inflammatory cytokines, increase of the Treg pool and modulate of innate immunity, while preserving the blood-brain barrier integrity and enhancing axonal and myelin preservation (51, 52, 54, 55, 104, 105, 107).

Here, we investigated whether physical exercise could further modulate innate immunity and promote recovery when applied therapeutically after the peak of disease in EAE. We show that low-intensity exercise at the onset of remission enhances microglia-node interaction. This correlates with increased pro-regenerative and homeostatic microglia, as well as improved clinical recovery, further supporting a beneficial role for reinforced neuron-microglia crosstalk at the onset of repair.

These effects may partly stem from exercise-driven increases in neuronal activity, supported by changes in firing patterns and c-FOS expression (108, 109). Microglia sense neuronal activity and adapt their behavior accordingly, including modulation of neural function (30, 34, 110–114), and we recently showed that, after demyelination, microglial phenotype is strongly shaped by activity patterns (38), which is particularly relevant to MS-associated axonal damage (115–119). As exercise also alters endocrine function, cortisol level and inflammation (120), it will be important to test whether selective neuronal activation alone can similarly promote repair in EAE by tuning neuro-glial crosstalk.

Our low-intensity exercise paradigm supports the use of tailored physiotherapy after relapse. Future work should determine how long these benefits last, and how exercise intensity and timing can be individualized to maximize recovery (121).

### The rate of microglia-node interaction correlates with repair in MS pathology

Finally, this work provides the first description of microglia-node of Ranvier interaction in pwMS. We analyzed shadow plaques, in comparison with normal appearing white matter (NAWM) and control tissue. Our observations in human samples mirror our findings in EAE, with reinforced interaction in remyelinating areas compared to NAWM from pwMS and control tissue, and a positive correlation between the frequency of microglia-node interaction and the extent of repair. Further characterization of axonal pathology, oligodendroglial lineage, and adaptive immune cell content within shadow plaques would help determine whether increased interaction is associated with lesions exhibiting reduced axonal damage or loss, higher oligodendrocyte density and/or maturation, and resolved inflammation.

Together, our data suggest that recruited peripheral immune cells can foster neuron-microglia crosstalk, driving microglial reprogramming and participate in a virtuous, pro-repair cycle. In contrast, weak microglia-neuron interaction associates with persisting pro-inflammatory states. This mirrors pwMS pathology, where active lesion edges are either conducive to remyelination or harbor persistent pro-inflammatory microglia at the rim of mixed active-inactive lesions. Overall, these findings point to a finely balanced microglial fate, shaped by competing signals, with profound impact on repair capacity.

## Material and Methods

### Animals

Mice care and use conformed to institutional guidelines and animal use regulations (UPMC, INSERM, French and European Community Council Directive 86/609/EEC). The following mouse strains were used: C57BL6/J (Janvier Labs), CX3CR1-GFP (gift from Prof S. Jung, Weizmann Institute of Science) and Synβ1Nav-mCherry, generated in collaboration with the transgenesis facility of Institut Curie, Paris. Briefly, β1Nav-mCherry sequence was extracted from custom-generated construct (Genecust) using EcoRI/SalI double digest and inserted, using the same restriction sites, downstream of the hSynapsin promoter sequence localized into pBluescript (Genecust custom sequence; pBS-EcoRV-hSyn-EcoRI-SalI-polyA). The resulting construct was used by the Institut Curie transgenesis facility to generate the Synb1Nav-mCherry line by classical pronuclei injection.

### EAE Induction and follow-up

#### Induction protocol

On day 0, 9-12-week old C57BL6/J female mice or CX3CR1-GFP/SynB1Nav-mCherry mice (both males and females) were sedated (Isoflurane 1.5%) and subcutaneously (s.c.) injected with 100 µL of MOG_35-55_ peptide (2mg/mL; MEVGWYRSPFSRVVHLYRNGK, Neosystem) emulsified with 100 µL complete Freund’s adjuvant (CFA, Sigma-Aldrich) containing 500 µg of mycobacterium tuberculosis (Coger), adapted from (61, 122). Control groups of matched littermates were injected with CFA/PBS only. On days 0 and 2, all mice received an intraperitoneal injection of pertussis toxin (PTX, Coger, 400 ng in PBS).

#### Scoring of disease severity

Mice were monitored daily for weight loss and clinical, and disease severity was scored as follows: 0, no detectable signs; 0.5, flaccid tail; 1, complete tail paralysis; 1.5, complete tail paralysis and hind limb inhibition; 2, partial hind limb paralysis; 2.5, unilateral complete hind limb paralysis or bilateral paresis; 3, complete bilateral hind limb paralysis; 3.5, complete hind limb paralysis with partial forelimb paralysis; 4, complete paralysis of forelimbs and hind limbs; and 5, death.

#### Study timepoints and groups

We focused on two key timepoints during EAE, the peak of disease severity and at the onset of remission. The peak was defined as 48h of sustained maximum severity score (SS) and was typically observed between 14 and 18 days post-induction (DPI 14-18). Remission was defined as sustained improvement of SS compared to the peak (DPI 19-23) (61). For this study, mice were divided in two subgroups according to their severity score at peak: the “Severe EAE” group (scores 3-4), and the “Moderate EAE” group (scores 2-2.5). In the severe EAE group, we investigated the correlation between microglia-node interaction and recovery following relapse using clinical assessment, behavioral testing, and analysis of *in vivo* fixed tissue analysis (see below) (n = 20, 5 independent experiments, 3-5 mice per experiment). In the moderate EAE group, we assessed the impact of physical activity on microglia-node of Ranvier interaction. Mice were matched for SS score at peak and then randomly assigned to either a “physical activity” or a “sedentary” group (see below) (n = 28, 3 independent experiments, 3-6 mice per group per experiment).

#### Behavioral testing

To assess hindlimb motor strength in mice with EAE, we used a Grip strength meter (BIOSEP). The meter was set up according to the manufacturer’s instructions, and mice were held by the tail and gently lowered towards the apparatus. Animals were allowed to grab the upper part of the metal grid and were then pulled downwards following the grid plane. The force applied to the grid when the grasp was released was recorded in grammes. As no prior training is required for this test, animals were assessed at baseline (prior to EAE induction), and then at two subsequent time points: during peak disease severity and at the onset of clinical remission. Grip strength measurements were performed in triplicate and averaged. The percentage of improvement from peak to remission was calculated for each individual mouse.

#### Physical Exercise Training

Adult female C57BL6/J mice were acclimated to handling by the experimenter for 1 week (5 min per animal per day) to reduce stress. Following acclimatation, mice underwent a 5-day treadmill habituation protocol (EXER-GAIT TREADMILL, Clever Sys. Inc) with incremental increase in duration, up to 10 continuous min at 5 cm/s). After EAE immunization, mice matched for disease severity (SS 2-2.5 at the peak) were randomly assigned to either a low-intensity exercise (LIE) or sedentary (SED) group (ratio 1:1). Mice in the LIE group completed 10-min sessions at 3 to 5 cm/s, three sessions per day spaced 2 h apart, for four consecutive days starting 48 h after peak disease. The speed range of 3 to 5 cm/sec was chosen to approximate the free-movement speed for mice with a motor score of 2 to 2.5, as measured in an open-field setting. Mice in the SED group were placed on a stationary treadmill for the same duration and number of sessions.

#### Automated gait analysis

Locomotor gait dynamics were assessed using the CatWalk automated gait analysis system at baseline (end of the habituation protocol, prior to EAE induction), pre-training (before the first training session), and post-training (after the final training session). Two videos of forced walking were analyzed per mouse at each timepoint, recorder by a camera incorporated in the EXER-GAIT treadmill setting. For a subset of mice, locomotion was further evaluated during free running using the RunWay system (Clever Sys. Inc), with the mean of two passages calculated for each timepoint.

#### Spinal glass window implantation for *in vivo* live imaging study

To perform *in vivo* imaging following immune-mediated demyelination, spinal glass window implantation was combined with EAE induction. The implantation protocol of a spinal cord window was adapted from (123, 124). Briefly, buprenorphine (0.2 mg/kg) and ketoprofen (5 mg/kg) were injected s.c. 15 min prior to anaesthesia. Adult CX3CR1-GFP/Synb1Nav-mCherry mice (both male and female) were anaesthetized with ketamine/xylazine (100 mg/kg; 10 mg/kg, intraperitoneally, i.p.) and supplemented hourly with half the ketamine dose alone (50 mg/kg, intramuscular injection, i.m.). Following a dorsal skin and muscle incision, the spine was immobilized with two spinal forks placed at T11 and L1 vertebrae. Two staples were fixed along the transverse processes of the vertebrae to support a reshaped paperclip, which was stabilized with glue (Cyanolite) and dental cement (Unifast Trad 250 mg/250 mg, GC Dental Products Corp). A dorsal laminectomy of T12 and T13 was performed using a high-speed drill with a carbide bur. The spinal cord was hydrated with a solution PBS/Dexamethasone 0.05% (Naquadem). Custom-made glass window, cut from a glass coverslip (Menzel-Glaser, 2mm width and 5mm length), was placed above silicon (Kwik-Sil, World Precision Instruments) directly applied onto the spinal cord, and the window was sealed with glue and dental cement. Animals were placed in a warming chamber (Vet tech solution LTD, HE011) and left to wake. Post-operatively, mice received ketoprofen (2.5 mg/kg, s.c.) on the day following surgery and every two-days for 1 week. EAE induction was performed as previously described four weeks after glass window implantation. To perform *in vivo* imaging following focal spinal cord demyelination, glass window implantation was combined with Lysophosphatidylcholine (LPC; Sigma-Aldrich L4129) spinal injection, following a similar to that described by (37). After laminectomy of T13 and exposure of the dorsal funiculus of the spine, 1 μL of LPC (10mg/mL) diluted in 9‰ NaCl was injected intraspinally using a glass capillary. A glass window was then implanted as described above, and mice were placed into a warming chamber until recovery.

### Longitudinal live-imaging study *in vivo*

Mice with EAE or LPC-induced lesions, implanted with a spinal cord window, were imaged at both the peak of disease severity and remission onset, or at the peak of demyelination (DPI 7) and at remyelination onset (DPI11), respectively. Mice were anesthetized as described previously, and anesthesia was maintained with additional Ketamin/Xylazin injections as needed throughout the experiment. Imaging was performed in a heated chamber at 34°C using an upright 2-photon microscope (Zeiss 710 NLO) equipped with a ×20 water objective (NA 1,0) and a Coherent Vision II laser. Focal lesions were identified by the visualization of microglial/macrophage aggregation and activation, and acquisitions were made in the lesion vicinity (<250µm). Z-stack series (1024 × 1024 pixels, 170 × 170 µm, 20-30 images per stack, z-step of 0.65 µm) were acquired at an excitation wavelength of 940 nm every 10 min for 1 h. Following imaging, mice were placed in a warming chamber until fully awake and then reimaged at 11 days post-LPC injection or at the onset of EAE remission. Branches of the central vein were used as a guide for localization on the tissue across sequential imaging sessions. Control mice (CFA/PTX) were imaged at the same timepoints as EAE mice. For analysis, nodes initially contacted by microglia were selected for acquisitions. We quantified the percentage of frames showing contact and the number of consecutive frames in which contact was maintained over 1h. Post-acquisition image processing was carried out semi-automatically using a custom-written script on ImageJ (NIH, Bethesda, Maryland). Briefly, for each timepoint, the z-stack was maximum-projected into a single image, filtered (Median filter 2.0), and background-subtracted (100 pixels in green). Images were then realigned using StackReg in ImageJ, and the realigned images were assembled into a movie representing each acquisition timepoint.

### Luminex assay

For cytokine and chemokine profiling, n = 12 severe EAE mice (severity score 3–3.5 at peak) were followed longitudinally until the onset of remission. Animals were euthanized by CO₂ inhalation, and the spinal cord was dissected. Cervical, upper thoracic, and lumbar segments were fresh-frozen for Luminex analysis, while the lower thoracic segment was post-fixed in 2% PFA for 2 h and subsequently cryoprotected as described above for histological processing.

For Luminex analysis, tissue homogenates were prepared in Cell Lysis Buffer (Invitrogen, EPX-99999-000) at a ratio of 500 μL per 100 mg of tissue, homogenized at 25 Hz for 1 min, and centrifuged at 16,000 × g for 10 min at 4 °C. Supernatants were collected, and protein concentration was determined using the DC Protein Assay Kit (Bio-Rad). Samples were normalized to 10 mg protein/mL in 1× PBS. For the assay, 25 μL of each sample was diluted 1:1 with Universal Assay Buffer and loaded in duplicate into the ProcartaPlex Mouse Immune Response Panel 64-plex (Invitrogen, EPX640-20064-901), according to the manufacturer’s instructions. A total of 62 cytokines/chemokines were reliably detected, with two analytes (IL-3, IL-5) below the detection threshold.

### Organotypic cultures of mouse cerebellar slices

Cerebellar slice cultures were made as previously described (125). Briefly, cerebella from P8-10 mice were dissected in ice cold Gey’s Balanced Salt Solution (G9779, Sigma-Aldrich) supplemented with D-Glucose (4.5 mg/mL; G8769-100ML, Sigma-Aldrich) and 1X penicillin-streptomycin (100 IU/mL, Thermo Fisher Scientific). Tissue was then cut into 250 μm parasagittal slices using a McIlwain tissue chopper and placed onto Millicell membranes (0.4 μm, Merck Millipore, 3-4 slices per animal) in culture composed of 50% Basal Medium Eagle (BME; 41010026, Thermo Fisher Scientific), 25% Hanks’ Balanced Salt Solution (14185-045, Thermo Fisher Scientific), and 25% heat-inactivated horse serum (26050088, Thermo Fisher Scientific) medium, supplemented with GlutaMax (2 mM; 35050038, Thermo Fisher Scientific), penicillin-streptomycin (100 IU/mL, Thermo Fisher Scientific), and D-Glucose (4.5 mg/mL; Sigma-Aldrich). Slices from a given animal were separated on two membranes: one used as a control and the other for treatment, enabling paired comparisons and minimizing inter-animal variability. Cultures were maintained at 37°C under 5% CO_2,_ and the medium changed every 2-3 days. Experiments were analyzed at 10 days *in vitro* (10 DIV, remyelination stage).

### Cytokine treatments of cerebellar slice cultures

To minimize inter-animal variability inherent to *ex vivo* culture systems, slices from each animal were cultured in parallel on two separate membranes. One membrane was used for treatment, while the other served as an internal untreated control, allowing for paired experimental comparisons. Demyelination was induced at 6 days *in vitro* (DIV6) by incubating all slices with 0.4 mg/mL Lysophosphatidylcholine (LPC; L4129, Sigma-Aldrich, Merck) for 15 h 30 min in fresh culture medium. At the end of the demyelination period (DIV8.5), slices from one membrane were incubated for 24 h with IL-13 (20 ng/mL; Sigma-Aldrich, I1896) and IL-4 (20 ng/mL; RnD Systems, 504RL005) diluted in culture medium, while slices on the paired membrane were incubated with culture medium alone. Slices were fixed at DIV 9.5.

### Fixation of cultured cerebellar slices

Cerebellar slices were fixed as previously described (125), using 4% paraformaldehyde (PFA; Electron Microscopy, Thermo Fisher Scientific) for 5 min followed by 1% PFA for 25 min at room temperature (RT), and then washed in 1× PBS (ET330-A; Euromedex). Subsequently, slices were incubated at −20 °C for 20 min in absolute ethanol (Sigma-Aldrich; for PLP staining) and washed again in 1× PBS. Slices were then blocked for 1 h in 1× PBS containing 5% normal goat serum (50-062Z; Thermo Fisher Scientific) and 0.3% Triton X-100 (9036-19-5; Sigma-Aldrich), followed by incubation with primary antibodies diluted in blocking solution overnight at RT. After washing in 1× PBS, slices were incubated for 3h at RT in the dark with secondary antibodies diluted in blocking solution. Finally, slices were washed in 1× PBS and mounted between a glass slide and a coverslip (VWR; 631-0138) with Fluoromount-G (0100-01; Southern Biotech).

#### Histology and Immunohistochemistry Antibodies

Primary antibodies: mouse IgG2b anti-AnkyrinG (clone N106/65; 1:75; NeuroMab), mouse IgG1 anti-Caspr (1:100; NeuroMab), rabbit anti-Caspr (1:300; Abcam), mouse IgG1 anti-Pan Nav (clone K58/35; 1:150; Sigma-Aldrich), rat anti-PLP (1:10; kindly provided by Dr. K. Ikenaka, Okasaki, Japan), mouse anti-MOG (1:5; Linington), rabbit IgG anti-Iba1 (1:500; Wako), mouse IgG1 anti-Iba (1:100; Abcam), rabbit IgG anti-TMEM119 (1:100; Sigma-Aldrich), mouse IgG1 anti-TMEM119 (1:100; Invitrogen), rabbit anti-P2Y12r (1:300; Alomone), mouse IgG1 anti-MHC-II (1:100; Dako), mouse IgG2a anti-iNOS (1:100; BD Biosciences), goat anti-IGF-1 (1:50; R&D System), mouse IgG1 anti-CD68 (1:200; Agilent), chicken anti-GFP (1:250; Millipore), chicken anti-mCherry (1:1000; Abcam), rabbit IgG anti-CCR2 (1:100; Abcam), rabbit IgG anti-CD3 (1:200; Agilent), mouse IgG1 neurofilament-H anti-SMI31 (1:1000; Biolegend-Covance), mouse IgG1 neurofilament-H anti-SMI32 (1:1000; Biolegend-Covance), rat anti-PDGFRα (1:100; BD Biosciences), mouse IgG1 NaBC1 (BCAS1, 1:100; Santa Cruz), mouse IgG2b anti-CC1 (1:200; Millipore), rabbit anti-Olig2 (1:200; Sigma-Aldrich). Secondary antibodies consisted of goat or donkey anti-chicken, anti-goat, anti-mouse (IgG1, IgG2a, IgG2b), anti-rabbit and anti-rat antibodies conjugated to Alexa Fluor 405, 488, 594, or 647 (1:500; Invitrogen).

#### Immunofluorescent stainings of mice tissue

For the *in vivo* fixed tissue analysis of EAE-induced mice and their controls, the lower thoracic spinal cord was extracted following intracardiac perfusion with 2% paraformaldehyde (PFA) and post-fixed for 60-180 min, washed in PBS, and incubated in 30% sucrose for 24h at 4°C. The tissue was cryoprotected in O.C.T (Tissue-Tek, Sakura), sectioned in longitudinally (20 µm) using a Leica CM3050S cryostat, and stored at −20°C until use. Tissues were first rehydrated in PBS. For myelin protein staining, tissues were pre-incubated in absolute ethanol at −20°C for 20 min. An additional post-fixation step with 4% PFA for 10 min was applied prior to blocking for some antibodies (CCR2, TMEM119, iNOS, AnkG). Following these pre-treatments, tissues were washed in 1× PBS. Tissues were then incubated in blocking buffer (5–10% goat or donkey serum albumin and 0.2% Triton X-100 in 1× PBS)) for 1 h, followed by incubation with primary antibodies in blocking buffer overnight at 4°C. After washing in PBS, tissues were incubated with secondary antibodies diluted in blocking buffer for 2h at room temperature in the dark. When applicable, tissues were immersed for 10 min in Hoechst solution (2.5 μg/mL; Euromedex). Finally, tissues were mounted under a coverslip (VWR) with Fluoromount (Southern Biotech) and stored at 4 °C.

### Human tissue

#### Tissue preparation

Human tissue was obtained after informed consent from the UK MS Society Tissue Bank at Imperial College, London, under ethical approval by the National Research Ethics Committee (08/MRE09/31), as snap-frozen blocks. Cryosections (16 µm; Leica CM1950) were prepared and stored at −80 °C until use.

#### Chromogenic immunohistochemistry

Human brain sections were fixed in 4% PFA for 10 min before elimination of endogenous peroxidase activity with 0.1% H_2_O_2_ (Sigma-Aldrich) in PBS for 20 min. Sections were blocked with 5% horse serum and 0.3% Triton X-100, and then incubated with primary antibodies diluted in blocking solution. Binding of biotinylated secondary antibodies (Vector Laboratories) was visualized using the avidin-biotin horseradish peroxidase complex (Dako, Biotin Blocking System) and 3,3’-diamino-benzidine (DAB) (Vector Laboratories) as substrate. In parallel, primary antibodies were also detected using the ABC-alkaline phosphatase detection system (Vector Laboratories) with Vector Blue as the substrate. Images were captured with a QICAM digital camera (QImaging Inc).

#### MS lesion characterization

Hemispheric brain tissue was obtained from n = 4 healthy donors and n = 6 MS patients. The first three slices of each block were used for tissue characterization according to the latest classification (Kuhlmann et al., 2017). Luxol fast blue (LFB)–periodic acid Schiff (PAS) staining was performed and consecutive sections were subjected to immunohistochemistry (IHC) for MBP and MHC-II to detect demyelination and assess myeloid cell density and distribution. We analyzed n = 4 NAWM areas (derived from blocks of non-lesioned MS tissue), and from a total of n = 30 white matter lesions in the cerebrum, we identified n = 10 remyelinating lesions. Patient and lesion data are summarized in Table 1.

#### Immunofluorescent staining of human tissue

For human post-mortem brain study, sections were rehydrated with PBS, fixed in 4% PFA for 5 min, then washed and incubated in blocking buffer (10% goat serum albumin, 0.2% Triton X-100 in 1× PBS) for 1 h. Sections were next incubated with primary antibodies in blocking buffer overnight at 4°C, washed in PBS, incubated with secondary antibodies diluted in blocking buffer for 2 h at room temperature in the dark. Finally, sections were mounted under a coverslip (VWR) with Fluoromount (Southern Biotech) and stored at 4°C. Neuroglial interactions at the node of Ranvier (AnkG, Nav) were assessed for homeostatic (P2Y12R) microglia, as well as resident (TMEM119) microglia. A fourth channel was used to extract lipofuscin signal during image analysis in ImageJ. Briefly, lipofuscin staining was removed from images by subtracting the “phantom” channel (without immunostaining) from each channel corresponding to nodal and microglial staining.

### Microscopy

#### Confocal microscopy

Confocal microscopy was performed using an inverted Leica SP8x microscope with a ×40 oil immersion objective (numerical aperture 1.30) and LasX software. Imaging of in vivo fixed tissues of EAE spinal cord: During the relapse, we focused on acute lesions identified by high densities of Iba1^+^ cells with “activated” morphology (reduced ramifications and/or hypertrophied cell body), high DAPI density, and loss of myelin staining. Microglia-node interaction was studied in the perilesional white matter (PPWM) within 250 µm of the lesion. During remission, lesions with partial resolution of inflammation, reappearance of ramified microglia, and presence of partially myelinated axons in the lesion were studied. Images were acquired with a ×40 oil objective (NA 1.30) and an optical zoom of 2, covering an area of 145.45 × 145.45 μm (0.0216 mm^2^). Five stacks of 1024 × 1024 pixels were collected from at least 3 different lesions per animal, with 10 sections per stack and a z-step of 0.30 μm. For quantification of cell and axonal densities, three stacks (1024 × 1024 pixels, ×40 oil objective NA 1.30, optical zoom of 1, 0.0844 mm^2^ area) were acquired on the section with the largest lesion.

Imaging of post-mortem human MS tissue: Following lesion characterization as described above, adjacent sections were used to quantify microglia-node interaction and microglial phenotypes. Images were acquired in NAWM and within remyelinated areas of shadow plaques. Following immunofluorescent staining, for each lesion type and subject, three stacks of images were acquired using a ×40 oil objective (NA 1.30) with optical zoom of 2, 10 images per stack, and a z-step of 0.30 μm.

#### Conventional fluorescent microscopy

To evaluate the extent of demyelination and inflammatory lesions in the spinal cord of EAE mice, we acquired mosaic images of entire thoracic spinal cord longitudinal sections (two to three sections/animal, selected based on the largest lesion areas). Images were obtained using an upright Axio imager microscope (Zeiss) with a 20× objective (numerical aperture 0.8, numerical zoom 1; 2048 × 2048 pixels, corresponding to a field area of 0.443 mm^2^ per image) and a Hamamatsu Flash 4.0 LT sCMOS camera. Images were stitched using Zen software (Zeiss). Demyelinated areas were defined by the absence of PLP staining and the presence of confluent Iba1^+^ cells. For each section, the proportion of demyelination was calculated as the total demyelinated area divided by the total section area. For each animal, values were averaged across two to three sections.

### Data Analysis

For quantification of microglia-node contacts in mouse *in vivo* fixed tissue, five images were analyzed per animal and per condition, from at least three different lesions, using ImageJ software. A contact was defined as at least one positive pixel for the microglial marker juxtaposed to at least one positive pixel for the nodal marker. The middle plan of each Z-series was used to calculate: (i) microglial density, (ii) nodal density, and (iii) the percentage of contacts, calculated as the number of nodes contacted divided by the total number of nodes. The mean percentage of contacts was then calculated per animal, and adjusted for microglial and nodal density per animal/condition.

To investigate the nature of contacting cells and assess whether resident microglial cells or recruited macrophages/activated microglia contact nodal structures, three images per animal/condition, (0.0216 mm^2^ area/image) were analyzed in two paradigms: (i) percentage of nodes contacted by double-positive TMEM119^+^/Iba1^+^ cells vs TMEM119-/Iba1^+^ cells in EAE-induced C57BL6/J mice; (ii) percentage of nodes contacted by GFP^+^/CCR2^-^ cells vs GFP^+^ or GFP^-^/CCR2^+^ cells in EAE-induced CX3CR1-GFP mice.

Densities of pro-inflammatory and pro-regenerative microglia/macrophage cells were calculated in the middle plane of each Z-series, three to five images per animal, in at least 3 different lesions (0.0216 mm^2^ area/image).

Median split was used to separate high vs low contacting animals in the Severe-EAE cohort (n=20 animals) and z scores were calculated for the whole cohort. The same approach was used to separate high vs low contacting animals in the subgroup (n=12) used for the Luminex analysis.

For the analysis of the secretome in the spinal cord tissue of Severe-EAE mice, cytokine and chemokine expression levels (pg/ml) were calculated (mean of two duplicates per animal). The percentage of nodes contacted was also calculated for each animal (in the lower thoracic spinal cord segment that was used for immunohistochemistry). Z scores were generated for each molecule using the mean expression and used to generate an unsupervised heatmap analysis. For the analysis of differential expressions of cytokines and chemokines between subgroups (clusters A, B and C as identified by the unsupervised heatmap analysis and the high- and low-contacting groups, defined as described above) mean expressions per group and z scores were calculated.

To calculate axonal coverage (neurofilament SMI31/32^+^ axons), three to five images per animal (0.0844 mm^2^/area) from the same lesioned areas used for microglia-node interaction quantification were analyzed. A maximum intensity orthogonal projection was generated, and the axonal channel was manually binarized and registered as an independent file. The area covered by SMI31/32^+^ axons was measured using ImageJ, and the mean of five areas per animal was calculated. The mean control axonal coverage was calculated from n = 3 control mice. For each EAE mouse, the mean axonal coverage was divided by the mean control coverage to generate a ratio.

For time-lapse imaging analysis, the definition of contact was the same as above. The percentage of timeframes with contact, as well as the maximum number of consecutive timeframes with and without contact, was then calculated.

### Statistical analysis

Images were anonymized using a custom script that assigned randomized filenames, thereby concealing experimental conditions. After completion of image analyses, files were de-anonymized to restore their original labels, and statistical analyses were then performed. All statistical analysis and data visualization were performed using R software v4.3.2 (R Development Core Team, 2023) and Prism (GraphPad, version 8.4.3). Group sizes and the statistical tests applied are indicated in the text or figure legends. Parametric tests were used for n ≥ 5 when distributions passed the Shapiro-Wilk normality test; otherwise, non-parametric tests were applied.

Log_2_ fold-change threshold of 0.5 and -log_10_(p value) threshold of 1.3 were used to produce a volcano plot of differential expression of cytokines/chemokines between groups. We used adjusted p value y 0.05 with, when applicable, the Benjamini–Hochberg procedure to control the false discovery rate (FDR). Significantly overexpressed cytokines/chemokines (n=19) were used to investigate network connections (STRING analysis). Gene Ontology (GO) enrichment analysis was performed to identify overrepresented biological processes among the selected proteins. The analysis was conducted using PANTHER.

Group comparisons were performed by fitting a multivariate linear regression model to the percentage of node of Ranvier contacted, with condition as the main factor, adjusted for total node of Ranvier number and cell density as covariates. Analysis of variance (ANOVA) was performed using the F-test (function ‘Anova’ in the car R package v3.1-2), followed by post hoc pairwise comparisons using the Tukey’s method for multiple comparisons (emmeans R package v1.8.9). Animals in remission were divided into two subgroups based on a median split: “Remission Low-contacting” and “Remission High-contacting”. Associations between the percentage of node of Ranvier contacted and biological/clinical variables were explored descriptively using partial least squares regression (PLSR). PLSR performs dimension reduction by constructing orthogonal components from manifest variables and handles multicollinearity via the Nonlinear Iterative Partial Least Squares (NIPALS) algorithm. Two PLSR models were fitted separately for each experimental measurement type, and a third model combined both types, each using two components, with the function ‘pls’ of the mixOmics R package (v6.26.0) (126). Correlation circles were used to evaluate the association of each variable with the PLSR components and to identify variables linked to node of Ranvier contact. Model performance was assessed on the training data using Pearson’s correlation coefficient between percentages of node of Ranvier contacted and the first and second PLSR components, as well as R² values from linear regression models of node of Ranvier contacted with the first two PLSR components.

Kaplan-Meier curves were used for survival analysis. Grubbs’ test was applied to identify potential outliers. All tests were two-sided, with significance set at p or adjusted p < 0.05.

Chat-GPT 5.1 was occasionally employed for targeted language editing and stylistic improvements of the manuscript. BioRender was used for graphical illustrations.

## Supporting information

Statistical table

## Contribution

AD, VP, GD and TR designed the study, VP, GD, TR, MSA and CP did the experiments, MC and AM participated in the Luminex study, VP and TR did the analysis, FXL participated in the statistical analysis, AD and VP and wrote the manuscript; GD, CL, RDP, FXL and BS participated in writing.

## Aknowledgments

We thank S. Jung for kindly providing us the CX3CR1-GFP mouse line and the London MS Tissue Bank for providing the post-mortem human tissues used for this study. We thank R. Maggliozzi and T. Kuhlmann for training TR and VP in immunohistochemical analysis of post-mortem human tissues. We thank ICM platforms implicated in this project: ICM.Quant, Claire Lovo and David Akbar, PhenoICMice, Histomics, ICV-3C, iGenSeq and DAC. We thank N. Sol-Foulon and B. Nait-Oumesmar for their critical reading of the manuscript. This work was supported by France SEP and ANR (NOREMY) fundings to AD, ICM and INSERM. VP was supported by Fondation Livio Glauser, Fondation Afenduli, Fondation Novartis and ECTRIMS Fellowships.

