## Supplementary figures and legends for "High neuron-microglia interaction at the node of Ranvier predicts recovery in an inflammatory model of Multiple Sclerosis"

### Supplementary figures and tables

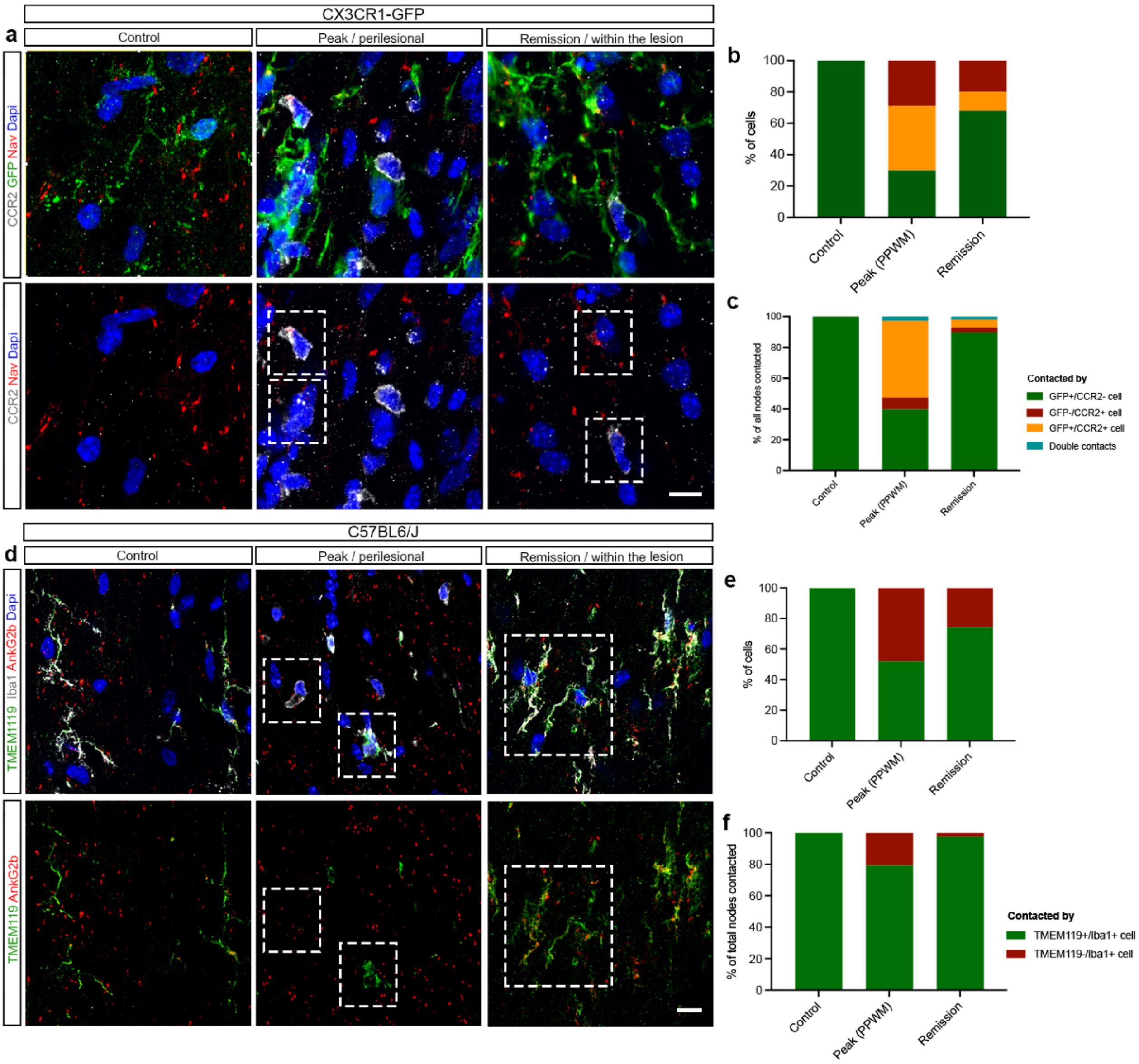


**Figure S1. Resident microglia cells are responsible for the majority of contacts at nodes at the onset of remission.**

**(a)** Immunostainings of thoracic spinal cord sections from EAE-induced CX3CR1-GFP mice showing microglia (GFP$^{+}$/CCR2$^{-}$, green) and monocyte subpopulations (infiltrating monocytes GFP$^{-}$/CCR2$^{+}$, grey, arrowheads; monocyte-derived macrophages GFP$^{+}$/CCR2$^{+}$, grey/green, arrows) interacting with nodes of Ranvier (Nav, red; microglia–node contacts indicated by white asterisks). **(b)** Mean cell density of microglia (green), monocytes (red), and MDMs (orange) during EAE progression (mean ±

SEM). **(c)** Percentage of nodes contacted by microglia (green), monocytes (red), and MDMs (orange) among all contacted nodes; a minority are contacted simultaneously by microglia and monocytes (blue) (n = 3–5 animals/condition). **(d)** Immunostaining of thoracic spinal cord sections from EAE-induced adult female C57BL/6J mice and matched controls showing resident microglia ($TMEM119^{+}$/$Iba1^{+}$, green) and activated microglia/MDMs ($TMEM119^{-}$/$Iba1^{+}$, grey) contacting nodes of Ranvier (AnkG2b, red; white asterisks). Mean cell densities for $TMEM119^{+}$/$Iba1^{+}$ (green) and $TMEM119^{-}$/$Iba1^{+}$ (red) are shown (mean ± SEM). **(e)** Distribution of $TMEM119^{+}$ vs $TMEM119^{-}$ cells among total $Iba1^{+}$ cells (% of $Iba1^{+}$ cells). **(f)** Percentage of nodes contacted by $TMEM119^{+}$/$Iba1^{+}$ vs $TMEM119^{-}$/$Iba1^{+}$ cells (n = 3–7 animals per lesion type). Scale bar: 10 μm.

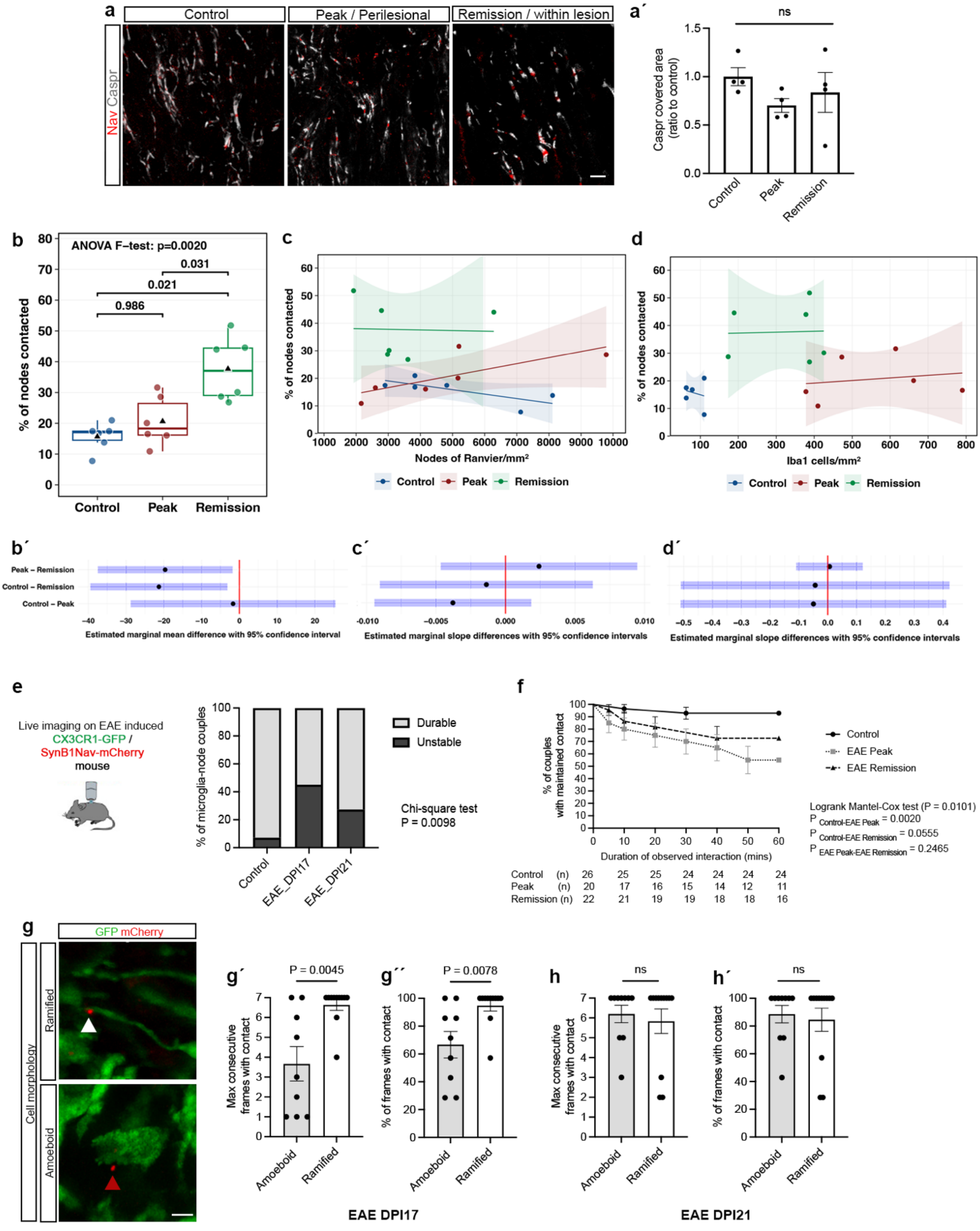
a
Control
Peak / Perilesional
Remission / within lesion
Nav Caspr
a´
ns
Caspr covered area (ratio to control)
1.5
1.0
0.5
0.0
Control
Peak
Remission
b
ANOVA F−test: p=0.0020
0.031
0.021
0.986
% of nodes contacted
Control
Peak
Remission
c
% of nodes contacted
Nodes of Ranvier/mm²
Control
Peak
Remission
d
% of nodes contacted
Iba1 cells/mm²
Control
Peak
Remission
b´
Peak – Remission
Control – Remission
Control – Peak
Estimated marginal mean difference with 95% confidence interval
c´
Estimated marginal slope differences with 95% confidence intervals
d´
Estimated marginal slope differences with 95% confidence intervals
e
Live imaging on EAE induced
CX3CR1-GFP /
SynB1Nav-mCherry
mouse
% of microglia-node couples
Durable
Unstable
Chi-square test
P = 0.0098
Control
EAE_DPI17
EAE_DPI21
f
% of couples with maintained contact
Duration of observed interaction (mins)
Control
EAE Peak
EAE Remission
Logrank Mantel-Cox test (P = 0.0101)
P Control-EAE Peak = 0.0020
P Control-EAE Remission = 0.0555
P EAE Peak-EAE Remission = 0.2465
Control (n) 26 25 25 24 24 24 24
Peak (n) 20 17 16 15 14 12 11
Remission (n) 22 21 19 19 18 18 16
g
GFP mCherry
Cell morphology
Ramified
Amoeboid
g´
P = 0.0045
Max consecutive frames with contact
g´´
P = 0.0078
% of frames with contact
h
ns
Max consecutive frames with contact
h´
ns
% of frames with contact
Amoeboid
Ramified
EAE DPI17
EAE DPI21

**Figure S2. Microglia-node interaction is independent of nodal and microglial density.**

**(a)** Images showing nodes of Ranvier (Nav staining, red), classified as mature nodes (flanked on both sides by paranodal Caspr, grey) in control tissue, heminodes (flanked on one side by Caspr), or pre-nodes (no Caspr segment) at EAE peak and at the onset of remission. **(a′)** Quantification of paranodal Caspr-covered area (ratio of Caspr area normalized by the mean Caspr area of n = 4 control mice) in the three conditions. **(b)** Adjustment for both nodal and Iba1$^+$ densities confirms a significant increase in microglia–node contacts during remission compared with control and peak ($F_{2,13} = 10.4$, $p = 0.002$). Black triangle represents the mean. **(c, d)** Scatterplots of percentage of nodes contacted vs total node number **(c)** or Iba1$^+$ cell density **(d)** for each condition, with linear regression lines and shaded 95% confidence intervals; no significant association detected ($F_{1,37} = 0.13$, $p = 0.72$ for **(c)**; $F_{1,12} = 0.11$, $p = 0.75$ for **(d)**). **(b′, c′, d′)** Pairwise comparisons of regression slopes between conditions, shown as mean differences in slopes with 95% confidence intervals (calculated using emmeans); intervals crossing zero (red line) indicate no significant difference. **(e)** In EAE, 35% of initially contacting microglia disengage from the nodes of Ranvier during remission vs 5% in control condition. **(f)** Kaplan-Meier survival curve for the duration of the observed interaction. The number of studied microglia/MDM-node pairs are indicated below the graph for each timepoint. **(g)** Representative images of GFP$^+$ cells (green) in contact with the nodes of Ranvier (mCherry, in red). Observed GFP$^+$ cells can be ramified (above) with small round cell body and elongated fine processes, or amoeboid (below) with enlarged cell body and maximum 3 proximal processes. White arrowhead indicates a contact between a GFP cell and a node of Ranvier and the red arrowhead indicates a detached GFP cell from the node. During peak, amoeboid cells detach from nodes **(g′-g′′)**, while cell morphology is not determinant for interaction stability during remission **(h′-h′′).** Statistical tests: Kruskal-Wallis test with Dunn's multiple comparisons test; ns, not significant in **(a′)**; F-test with Tukey's post-hoc pairwise comparisons, adjusted for total node of Ranvier number and Iba1$^+$ cell density, in (b-d); Chi-square test in **(e)**; Mann-Whitney test in **(g-h)**. Scale bar: **(a)** 10 µm, **(g)** 5 µm.

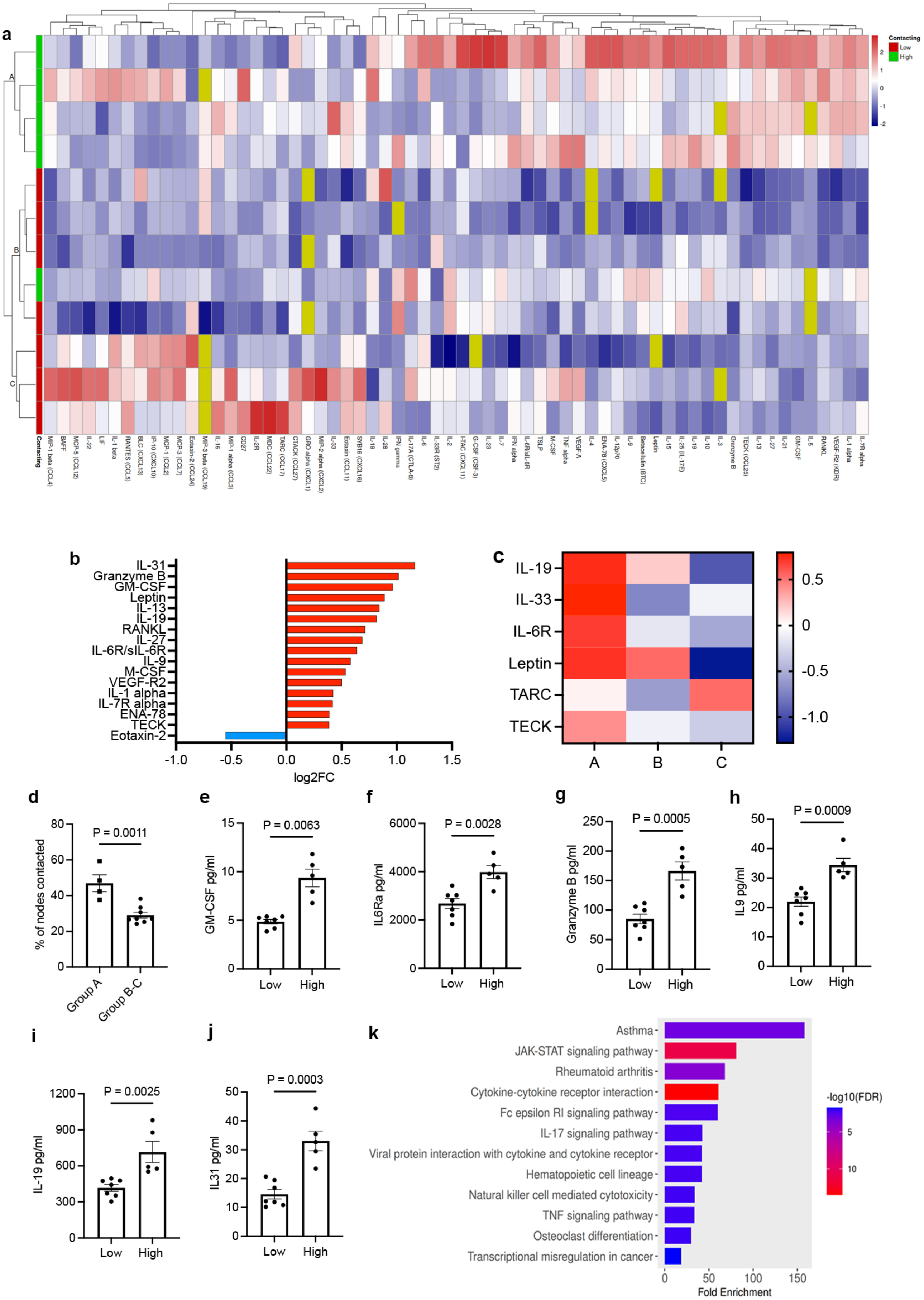
a
Contacting
Low
High
A
B
C
b
IL-31
Granzyme B
GM-CSF
Leptin
IL-13
IL-19
RANKL
IL-27
IL-6R/sIL-6R
IL-9
M-CSF
VEGF-R2
IL-1 alpha
IL-7R alpha
ENA-78
TECK
Eotaxin-2
-1.0
-0.5
0.0
0.5
1.0
1.5
log2FC
c
IL-19
IL-33
IL-6R
Leptin
TARC
TECK
A
B
C
0.5
0
-0.5
-1.0
d
P = 0.0011
% of nodes contacted
Group A
Group B-C
e
P = 0.0063
GM-CSF pg/ml
Low
High
f
P = 0.0028
IL6Ra pg/ml
Low
High
g
P = 0.0005
Granzyme B pg/ml
Low
High
h
P = 0.0009
IL9 pg/ml
Low
High
i
P = 0.0025
IL-19 pg/ml
Low
High
j
P = 0.0003
IL31 pg/ml
Low
High
k
Asthma
JAK-STAT signaling pathway
Rheumatoid arthritis
Cytokine-cytokine receptor interaction
Fc epsilon RI signaling pathway
IL-17 signaling pathway
Viral protein interaction with cytokine and cytokine receptor
Hematopoietic cell lineage
Natural killer cell mediated cytotoxicity
TNF signaling pathway
Osteoclast differentiation
Transcriptional misregulation in cancer
Fold Enrichment
-log10(FDR)

**Figure S3. Molecular candidates involved in interindividual heterogeneity of microglia-node interactions.**

**(a)** Unsupervised heatmap of Luminex-based multiplex analysis from n = 12 Severe EAE animals, showing relative expression (z-scores) of all molecules analyzed across the tested cohort. **(b)** Differentially expressed molecules ($log_2$ fold change, $log_2FC$) in group A compare with groups B and C. **(c)** Significant changes in expression ($log_2FC$) between groups A-B, B-C, and C-A. **(d)** The percentage of nodes contacted was significantly higher in group A (mean ± SEM). **(e-j)** Concentrations of selected cytokines/chemokines in low- vs high-contacting mice: **(e)** GM-CSF, **(f)** IL-6Ra, **(g)** granzyme B, **(h)** IL-9, **(i)** IL-19, and **(j)** IL-31 (mean ± SEM). **(k)** KEGG pathway analysis showing main pathways associated with the 19 upregulated molecules in high-contacting mice; bar color scale reflects $-log_{10}$ fold change. Statistical analysis: t-test with Welch's correction when applicable in **(d-j)**.

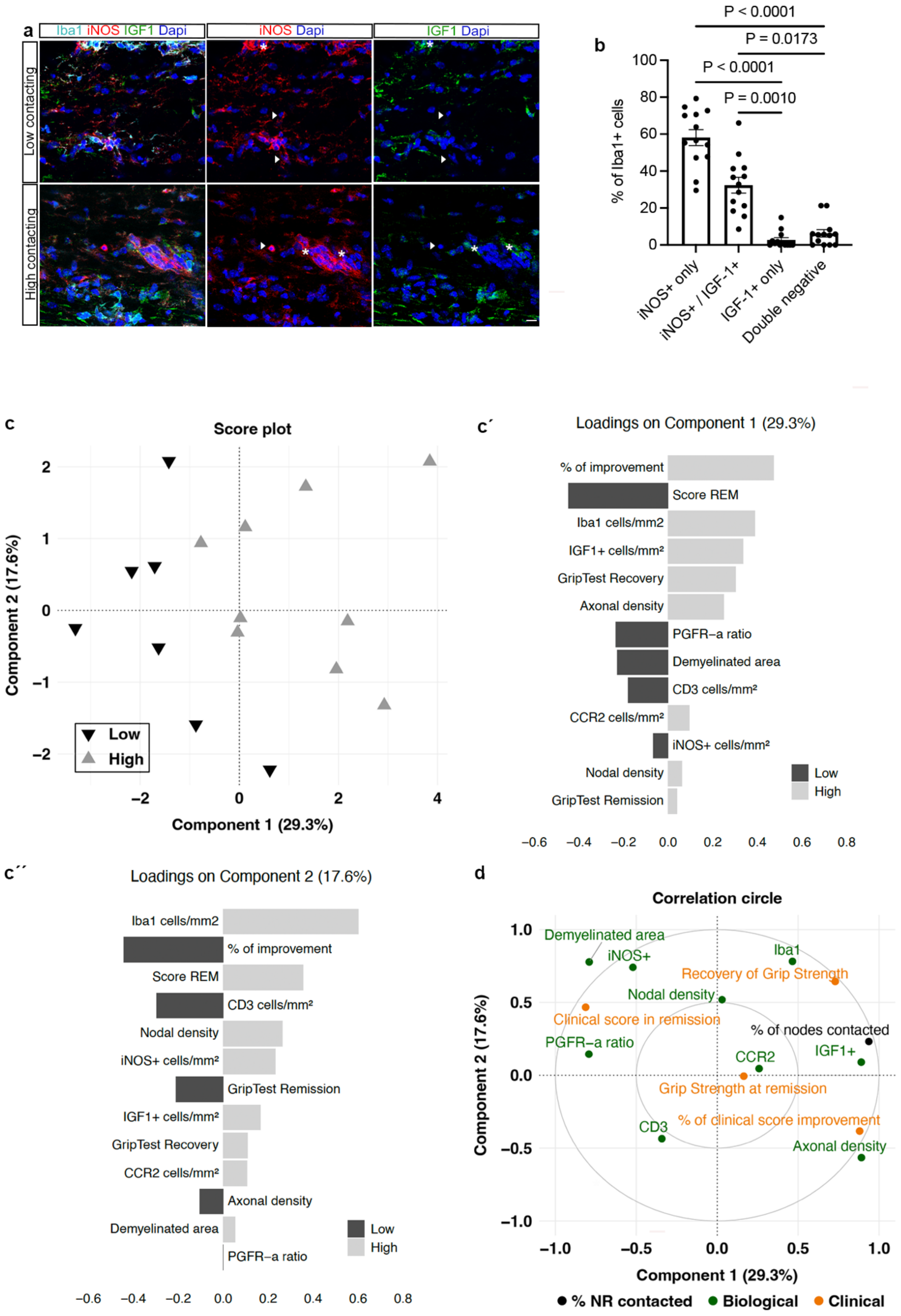
a
Iba1 iNOS IGF1 Dapi
iNOS Dapi
IGF1 Dapi
Low contacting
High contacting
b
% of Iba1+ cells
P < 0.0001
P = 0.0173
P < 0.0001
P = 0.0010
iNOS+ only
iNOS+ / IGF-1+
IGF-1+ only
Double negative
c
Score plot
Component 1 (29.3%)
Component 2 (17.6%)
Low
High
c´
Loadings on Component 1 (29.3%)
% of improvement
Score REM
Iba1 cells/mm2
IGF1+ cells/mm²
GripTest Recovery
Axonal density
PGFR–a ratio
Demyelinated area
CD3 cells/mm²
CCR2 cells/mm²
iNOS+ cells/mm²
Nodal density
GripTest Remission
Low
High
c´´
Loadings on Component 2 (17.6%)
Iba1 cells/mm2
% of improvement
Score REM
CD3 cells/mm²
Nodal density
iNOS+ cells/mm²
GripTest Remission
IGF1+ cells/mm²
GripTest Recovery
CCR2 cells/mm²
Axonal density
Demyelinated area
PGFR–a ratio
Low
High
d
Correlation circle
Demyelinated area
iNOS+
Iba1
Recovery of Grip Strength
Nodal density
Clinical score in remission
% of nodes contacted
PGFR–a ratio
CCR2
IGF1+
Grip Strength at remission
CD3
% of clinical score improvement
Axonal density
Component 1 (29.3%)
Component 2 (17.6%)
% NR contacted
Biological
Clinical

**Figure S4. Multi-modal integration of clinical and biological data suggests that high microglia-node interaction is associated with improved repair.**

**(a)** Co-staining to assess iNOS (red) and IGF-1 (green) expression in $Iba1^+$ cells (light blue) during clinical remission in Severe EAE. **(b)** Quantification of distribution of $iNOS^+$ only, IGF-$1^+$ only, double-positive, and double-negative cells among all $Iba1^+$ cells (mean ± SEM; n = 13 animals per condition). Each dot represents the mean of 5 areas analyzed per mouse and a minimum of 20 $Iba1^+$ cells per animal. Statistical analysis: Kruskal-Wallis test with Dunn's post hoc comparisons. **(c)** Partial least squares regression (PLSR) modeling (function 'pls' from mixOmics R package, v6.26.0) integrating all clinical and biological parameters studied in the Severe EAE subgroup. Frequency of microglia-node contacts was strongly associated with Component 1 (r = 0.91, 95% CI 0.76-0.97, p = 9.8e-7) but not with Component 2 (r = 0.33, 95% CI -0.20-0.71, p = 0.21). Component 1 explained 29.3% of variability, separating low- vs high-contacting mice and identifying highlighting discriminating variables. The loadings on component 1 and 2 are shown in **(c′)** and **(c′′)** respectively. **(d)** Correlation circle of the two-component PLSR model showing both biological (green) and clinical (orange) modalities, with % of nodes contacted added in black. Statistically significant variables are located above or below ±0.5 along the horizontal axis (Component 1). Correlations with microglia–node interaction are positive when above 0.5 and negative when below -0.5. IGF-$1^+$ cell density, axonal density, and clinical recovery were strongly associated with increased interaction. Scale bars: (a) 10 µm.

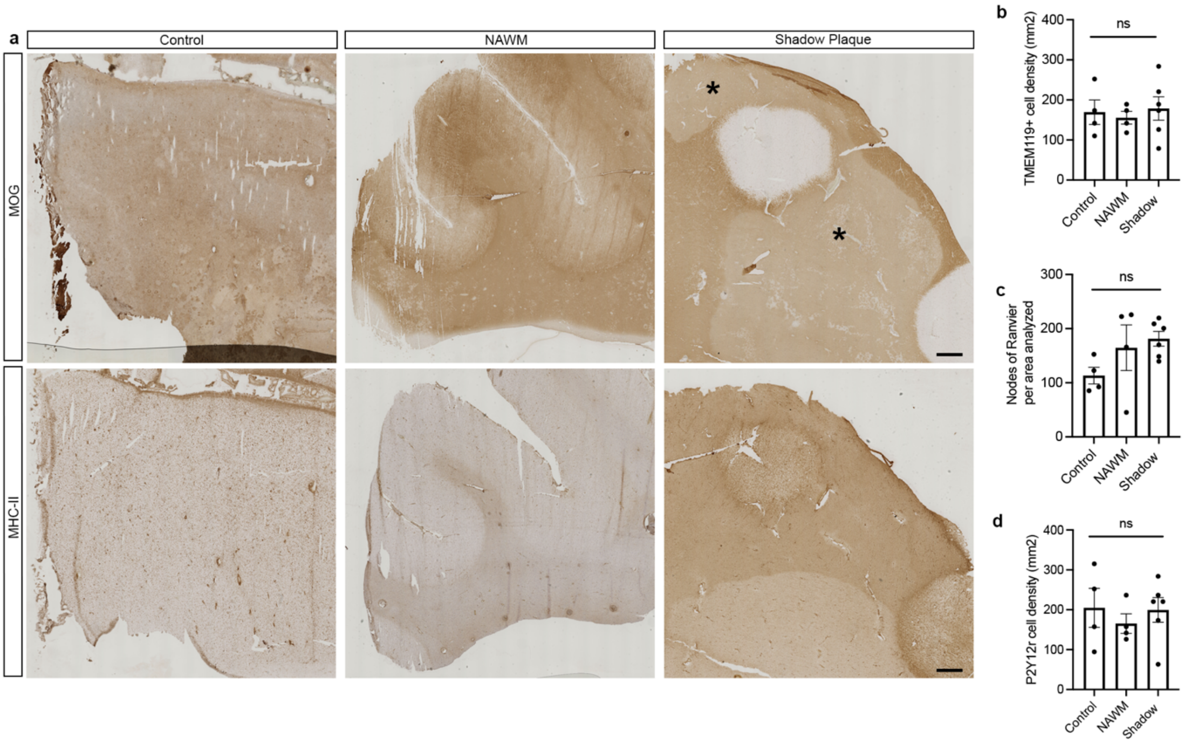


**Figure S5. MS tissue characterization.**

**(a)** Immunohistochemistry staining for MOG and MHC-II in the same tissue, in control and MS post-mortem hemispheric brain tissue. Control tissue shows normally myelinated cortex and white matter. MS tissue exhibits different disease activities: normal-appearing white matter (NAWM) with no myelin loss; and shadow plaques with pale MOG staining (indicated with an asterisk). **(b)** TMEM119$^+$ cell density/mm$^2$, **(c)** nodal density/mm$^2$ in the same areas analyzed for TMEM119$^+$ cells (n = 4-6 samples per condition), and **(d)** P2Y12R$^+$ cell density/mm$^2$. Kruskal-Wallis test; ns, not significant. Scale bar: 500 μm.

| *ID* | Sex | Age | Death to tissue preservation (hours) | Disease course | Disease duration (years) | Lesion types |
|---|---|---|---|---|---|---|
| *MS160* | F | 44 | 18h | SP | 16 | CAI, I |
| *MS166* | F | 52 | 7h | SP | 36 | N, S |
| *MS179* | F | 70 | 20h | SP | >21 | N, CAI, S |
| *MS180* | F | 44 | 9h | SP | >10 | A |
| *MS187* | F | 57 | 13h | SP | >27 | N, I, S |
| *MS207* | F | 46 | 10h | SP | 25 | N, CAI, I, S |
| *MS317* | F | 48 | 21h | SP | 29 | N, A, CAI, S |
| *C028* | F | 60 | 13h | N/A | N/A | N/A |
| *C054* | M | 66 | 16h | N/A | N/A | N/A |
| *C064* | F | 63 | 21h | N/A | N/A | N/A |
| *C074* | F | 84 | 22h | N/A | N/A | N/A |

**Supplementary Table 1. Human Brain Tissue samples.**

Abbreviations: F/M: Female/Male; N: NAWM (Normal Appearing White Matter); A: Active lesion; I: Inactive lesion; CAI: Chronic Active/Inactive lesion; S: Shadow plaque; N/A, Not Applicable.
